# Electro-Calcium uncoupling precedes neurodegeneration in Alzheimer’s disease

**DOI:** 10.64898/2026.01.26.701803

**Authors:** Hwoi Chan Kwon, Adrian Eiden, Joshua Li, Martin MacKinnon, Joshua B. Garfinkel, Sarah M. Hooper, Yanling Liu, Mark T. Nelson, Alan Koretsky, Amreen Mughal

**Author notes:** Co-first authors (Hwoi Chan Kwon, Adrian Eiden). Corresponding author Correspondence to Dr. Amreen Mughal, Full address: 35 Convent Drive, Bldg 35A Room 3D-933, Bethesda, MD 20892.

## Abstract

Alzheimer’s disease (AD) is categorized as a neurodegenerative disease, but there is a growing recognition of the vascular components in AD pathophysiology. Reduction in cerebral perfusion is routinely observed in AD patients and preclinical models prior to overt clinical symptoms. However, there is a limited mechanistic understanding of the early neurovascular deficits in AD, and how these may ultimately contribute to pathology. Here, we investigated the mechanisms of early neurovascular dysfunction in AD by using 3-month-old 5xFAD mice, a familial mouse model of AD. Functional hyperemia—the increase in cerebral blood flow (CBF) in response to neuronal activity—is driven by inward rectifier K^+^ (K_ir_2.1)-mediated hyperpolarizing (electrical) signals and Ca^2+^-dependent nitric oxide production within the capillary endothelial cells (cECs). Electrical and Ca^2+^ signals are tightly coupled through cECs membrane potential, referred to as Electro-Calcium (E-Ca) coupling. We hypothesize that E-Ca uncoupling contributes to impaired functional hyperemia in 5xFAD mice and that these neurovascular deficits precede the neurodegeneration and cognitive decline. At three months of age, 5xFAD mice did not exhibit any impairment in spatial learning and memory, or neuronal density. However, whisker stimulation-induced functional hyperemia was significantly reduced in 5xFAD mice compared to controls. Functional hyperemia exhibited a bimodal response in controls—consisting of fast and slow phases—with the slow phase being significantly reduced in 5xFAD mice. To identify mechanisms underlying these deficits, we measured cortical neuronal and endothelial Ca^2+^ activity using *in-vivo* imaging. Neuronal Ca^2+^ activity was comparable between controls and 5xFAD mice, while cECs Ca^2+^ activity was significantly reduced in 5xFAD mice. Moreover, K_ir_2.1 channel blocker, barium (100 μM) significantly suppressed cECs Ca^2+^ activity in controls, but not in 5xFAD mice, consistent with crippled E-Ca coupling. Despite these vascular functional impairments, capillary density was preserved in 5xFAD mice. TRPV4 channels are one of the major Ca^2+^ entry pathways in cECs and potentiate E-Ca coupling. cECs TRPV4 current density was significantly reduced in 5xFAD mice while K_ir_2.1 current density was unchanged, indicating that impaired TRPV4 function underlies the E-Ca uncoupling. In summary, early E-Ca uncoupling leads to impaired functional hyperemia in 5xFAD mice and may contribute to later neuronal and cognitive decline.

## 1. INTRODUCTION

The brain is one of the most metabolically “expensive” organs and is second only to the liver in absolute energy expenditure^1–4^. Neuronal action potentials and synaptic transmission consume a large amount of energy ^5,6^. The network of the brain vasculature senses and meets this energy demand through processes known as neurovascular coupling (NVC) ^7,8^. These NVC mechanisms are also critical in the clearance of metabolic waste (including K^+^, glutamate, amyloid beta and p-tau protein) among many other brain functions ^9^. Reduction in cerebral blood flow (CBF) is reported as an early change in AD patients^10^. However, most of the preclinical mechanistic studies are performed during symptomatic (neurodegenerative) phase of AD ^11–13^, leaving a gap in the knowledge about the NVC impairment during presymptomatic (early) phase of AD.

Reduced CBF and glucose metabolism are reported in human AD patients ^14–18^ and animal models with AD pathology ^11,12,19–21^. Moreover, CBF reduction manifests before the structural changes (grey/white matter atrophy) and clinical onset of dementia in AD patients^15,18,22^. This early reduction in CBF may be central for driving the neurodegeneration and cognitive decline in AD due to impaired energy supply and accumulation of metabolic wastes in the brain. With the number of dementia cases projected to reach ∼14 million by 2060^23^, identification and development of early therapeutic targets and interventions are critical^24^. Therefore, it is imperative to study early NVC deficits that are contributing to CBF impairment in AD.

Capillaries are the smallest blood vessels (in size; 3-8 µm) and are tightly interwoven throughout the brain parenchyma ^2,8,25^. The close proximity of capillaries to neuronal cells (∼8 µm) make them ideal for meeting neural energy demand and clearing metabolic waste ^7,8^. Given this proximity, any changes in capillary functions can be detrimental to blood flow regulation and neuro-vascular communication. We and others have identified NVC mechanisms that rely on capillary endothelial cells (cECs); namely electrical and Ca^2+^ signaling. Electrical (hyperpolarizing) signals are generated in cECs by virtue of K^+^ channels in response to modest elevation in extracellular K^+^ and adenosine with neuronal activity ^26–28^. Ca^2+^ signals in cECs are dependent on neuronal activity derived ligands that activate G_q_-protein coupled receptor (G_q_PCR) signaling and non-selective cation channels (e.g., TRPV4, TRPA1 and Piezo1) in cECs ^29–31^. The electrical (hyperpolarizing) signals initiate Ca^2+^ signals through an increase in Ca^2+^ entry caused by the membrane potential hyperpolarization and is referred to as Electro-Calcium (E-Ca) coupling^32^. Considering the crucial role of E-Ca coupling in regulation of CBF, we hypothesized that impairment in E-Ca coupling leads to early (presymptomatic phase) reduction in CBF in AD and these changes precede the neurodegeneration.

We used a familial genetic model of AD (3-month-old 5xFAD mice), with a combination of *in-vivo* and *ex-vivo* imaging approaches along with behavioral testing and electrophysiology. We show that in 3-month-old 5xFAD mice: (1) Whisker stimulation-induced functional hyperemia is impaired, whereas resting-state perfusion is not affected, (2) cECs Ca^2+^ signaling is significantly impaired due to reduced TRPV4 activity, resulting in E-Ca uncoupling, (3) cortical neuronal density and neuronal Ca^2+^ activity remain unchanged, and (4) these early NVC deficits precede the long-term spatial learning and memory impairment. These findings provide novel molecular insights about the reduced functional hyperemia in AD and may guide therapeutic inventions in rescuing CBF at the early stages of AD.

## 2. METHODS

### 2.1 Animals

All experimental procedures were approved by the NINDS Institutional Animal Care and Use Committee, in accordance with NIH guidelines for the use of animals. 5xFAD (Jax # 034848, C57bl6 background), *Cdh5*-GCaMP8 and EC Kir2.1-KO mice were group-housed on a 12-hour light:dark cycle with environmental enrichment and free access to food and water. 5xFAD and *Cdh5*-GCaMP8 were cross bred to permit visualization of endothelial Ca^2+^ signals in the presence of the 5xFAD mutations. EC K_ir_2.1-KO mice were generated by crossing mice with loxP-flanked K_ir_2.1 sites (Kir2.1fl/fl) with mice expressing inducible Cre recombinase under the transcriptional control of Cdh5 (Cdh5-CreERT2)^32^. For induction of EC Kir2.1-KO mice, offspring were treated with tamoxifen by intraperitoneal injection (80 mg/kg/d) for 5 consecutive days, followed by a minimum 7-day washout period before experimental interventions. Adult male mice (12±1 weeks) of and age-matched controls were used for all experiments.

#### Chemicals

Texas red dextran (70 kDa) was obtained from Invitrogen (USA). Elastase, neutral protease, and collagenase type 1 were procured from Worthington Biochemical (USA). Unless otherwise noted, all other chemicals were obtained from Sigma-Aldrich (USA) or VWR (USA).

### 2.2 Behavioral testing

The experiments were conducted in a quiet and low-lit environment to minimize animal stress. All behavioral equipment was cleaned with 70% ethanol between trials to eliminate effects of odors on behavior. Overhead videos cameras were used to capture animal behavior for subsequent analysis.

#### Open field test (OFT)

This test is used to identify any differences in anxiety or motor function between the 5xFAD animals and controls. The animals were placed in a 46 x 46 x 50 cm box and allowed to freely explore for ten minutes. The center area measured 23 x 23 cm and time spent in this area was recorded based on the location of the center of the animal as identified by TopScan analysis software, which was used for analysis of behavior sessions.

#### Y-maze

This test was used to assess spatial learning and memory. Test sessions began when an animal was placed into one arm of the opaque acrylic maze that consisted of three arms, each 21 cm long and 7 cm wide with 15 cm high walls, angled 120 degrees apart. For 8 minutes animals explored the maze. A spontaneous alteration occurred when a mouse visited the arm least recently visited, resulting in a sequential visit to each arm. The arm entries and alterations were calculated using the AnyMaze mouse tracking and analysis software. The percent spontaneous alterations were calculated as [(number of alterations)/ (total arm entries – 2)] x100. The first two arm entries were excluded as a potential spontaneous alteration.

#### Novel Object Recognition (NOR)

This test was used to assess long term memory. Mice were allowed to habituate to the behavioral chambers (46 x 46 x 50 cm boxes) for 10 minutes. Then mice were trained for 10 minutes with two identical objects in opposite corners of the chamber. After 24 hours animals were returned to the behavioral chamber where one of the familiar objects from the training phase was replaced with a new novel object of different shape and texture. Animals were considered to be exploring an object when the nose of the animal was both within 50 mm of the object center and pointed at the object center (head position <180° from object center). Animal tracking and analysis were performed using TopScan software. The discrimination index was calculated as (novel object exploration time / total exploration time) x 100.

### 2.3 Functional Ultrasound

Functional ultrasound (fUS) imaging was performed in skull-thinned mice^33^. Briefly, mice were anesthetized using isoflurane (5% induction, 1.5-2% maintenance). The scalp and periosteum were cut away, and the surgical area was thinned to an approximate thickness of 100-150 µm until the skull was evenly translucent. After the surgery, the thinned skulled area was covered with silicone elastomer (Kwik-Sil, WPI) for protection during recovery and meloxicam (5 mg/kg, s.c.) was administered subcutaneously for post-operative pain management. The mice were given 3-4 days to recover before performing imaging.

Iconeus One system (Iconeus, Paris, France) equipped with linear ultrasound probe (IcoPrime-15MHz, Iconeus) mounted on a linear motorized stage was used for fUS. The probe consists of 128 piezoelectric elements with a 15 MHz central frequency and 0.1mm pitch, with a FOV of 14×19×0.4 mm^3^ with an in-plane spatial resolution of 100 µm^2^. The imaging sequence and real-time doppler reconstruction were obtained using Icoscan software (Iconeus, France). The power doppler (PD) image was formed from ultrasound sequence transmitting 11 tilted planar waves from −10° to 10° in 2° increment at a frequency of 5.5 kHz, and the compound image was formed at 500 Hz framerate. The compound images were combined into a block, and each block was filtered using singular value decomposition clutter filtering to separate tissue signals from blood signal. These processed blocks produced the final PD image at a sampling rate of 2.5 Hz ^34,35^.

#### 2.3.1 Data acquisition

On the day of imaging, the skull-thinned mouse was anesthetized with 100 mg/kg ketamine and 10 mg/kg xylazine cocktail and head fixed to a stereotaxic frame. The area between the skull and ultrasound probe was filled with ultrasound gel (Aquasonic). A 3D angiogram of the image was taken using Icoscan, scanning anterior from lambda in 0.1 mm step for 5 mm. The parameters were chosen to include the entire barrel field of the somatosensory cortex. Icostudio (Iconeus) was used to align the angiogram to the Allen mouse brain atlas, and the aligned structure was used to identify the desired coronal slice over the somatosensory cortex barrel field (SSC-BF). The imaging coordinates were copied and moved to Icoscan and probe was aligned to the area of interest.

The total duration of recording (240 sec) consisted of 30 sec whisker stimulation (WS) period flanked on each side by 60 sec baseline period. The recording was triggered using custom controller box and software sending 5V transistor-transistor logic (TTL) signal to Iconeus for time-synchronization. Mouse whiskers were stimulated with 20 psi breathable air puff (70% nitrogen and 30% oxygen) given by a pulse width modulation (PWM) at a frequency of 20 Hz and 50% duty cycle through a pneumatic solenoid with a custom designed slit. PD signals were collected from both contralateral and ipsilateral cortex, where ipsilateral signals were used as the within experiment control.

#### 2.3.2 Data analysis

##### 2.3.2.1. Preprocessing

PD signals from SSC-BF were averaged along the spatial dimension. The time course of averaged PD signal was further preprocessed using a custom Python code before analysis. Preprocessing involved the following: the time course was first detrended by fitting a linear line through the baseline period as defined. The detrended data was lowpass filtered using Butterworth at approximately 1Hz based on fast Fourier transform of the time series data that retained 99% of the cumulative power of the PD. The average of first baseline period (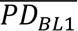) was used to normalize the trace to obtain a relative CBV (rCBV).

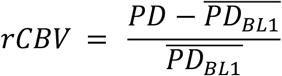

The average of the steady state rCBV during the first stimulation period is used for comparison.

##### 2.3.2.2. Activation map

The activation map was generated from the 4D images exported from Icostudio as Nifti image and using custom Python script. The y-dimension was collapsed for analysis purpose as it has a dimension of one. Then the images were first detrended pixel-wise and using a response model based on whisker stimulation parameter, pixel-wise Pearson product-moment correlation coefficient (r) map was generated ^36^.

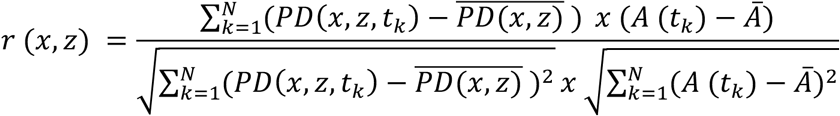

where N is the number of time points, PD is power doppler pixel time series, and A is the response model. The resulting map was masked with thresholding (σ = 3).

##### 2.3.2.3. Kinetic analysis

The two phases of activation were fit with a single-exponential rise function:

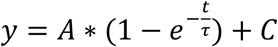

where y is the normalized rCBV, A is the amplitude of the fit, t is time, τ is time constant, and C is the offset. Where rise time constant (τ_rise_) is the time required to reach 63% of maximum response.

The drop in amplitude (A_drop_) was the difference between C_slow_ and A_fast_, while the delay between fast and slow phase (ΔT_drop_) was the difference between time when fast phase ended (T_fast-end_) and time when slow phase started (T_slow-start_).

Meanwhile, the decay was fit with single-exponential decay function:

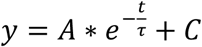

where the variables are same as above, except the decay time constant (τ_dec_) is the time to decay to 37% of the peak amplitude.

### 2.4 Functional and structural MRI

All MRI experiments were performed on an 11.7 T system (30 cm horizontal bore; Magnex Scientific, Oxford, UK) equipped with an AV NEO console (Bruker BioSpin, Ettlingen, Germany) and a BFG 240/120 gradient insert (Resonance Research Inc., Billerica, MA) with a rise time of 150 µs and a maximum gradient amplitude of 942 mT/m, operated with ParaVision 360.3.5 software. Data were acquired with an 86 mm volume coil for transmission and a 4-channel CryoProbe for reception (Bruker BioSpin, Ettlingen, Germany).

#### 2.4.1 Image Acquisition

A 2D whole-brain spin-echo based multi slice multi echo (MSME) acquisition was performed for anatomical localization for cerebral blood flow (CBF) measurements and subsequent registration to an anatomical atlas, with repetition time (TR): 1000 ms, effective echo times: 10, 20, 30 and 40 ms, Flip Angle: 180°, Field of View (FOV): 19.2 x 19.2 mm, Matrix Size: 192 x 192, Receiver Bandwidth: 50,000 Hz, slices: 20, Slice Thickness: 1 mm. With a nominal in plane resolution of the MSME at 100 µm.

The first labeling sequence used a short labeling duration of 200 ms to measure the degree of labeling and thereby assess labeling efficiency. A flow-compensated fast low-angle shot (fc-FLASH) acquisition was used for measurement of inversion efficiency, with the labeling plane set 2 mm caudal to the imaging plane at the bifurcation of carotid arteries with TR: 250 ms, Echo Time (TE): 3.4, FOV: 19.2 x 19.2 mm, Matrix Size: 128 x 128, Receiver Bandwidth: 50,000 Hz, Label gradient amplitude: 10 mT/m, Labeling Duration: 200 ms, Post Labeling Delay: 10 ms, B_1_: 12 µT, Slice Thickness: 1 mm, Averages: 2.

CBF was measured with a pseudo-continuous ASL (pCASL) sequence with an echo-planar-imaging (EPI) readout with parameters optimized in preliminary experiments (data not shown) with TR: 4,100 ms, TE: 13.84, FOV: 19.2 x 19.2 mm, Matrix Size: 64 x 64, Receiver Bandwidth: 357,143 Hz, Label gradient amplitude:10 mT/m, Labeling Duration: 3,000 ms, Post Labeling Delay: 300 ms, B_1_: 12 µT, Slice Thickness: 1 mm, Repetitions: 30, resulting in an acquisition duration of 3 m 40 s. The imaging slice was positioned to encompass the barrel field region of somatosensory cortex and labeling slice was positioned identically as the pCASL acquisition. For CBF quantification, the T1 of tissue was measured with a rapid acquisition with a saturation recovery refocused echoes sequence with based variable repetition time saturation recovery sequence (RARE-VTR) with Repetition Times (TRs): 12000, 8000, 3000, 1499, 800, 500, 395 and 100 ms, TE: 7.46 ms, FOV: 19.2 x 19.2 mm, Matrix Size: 64 x 64, Receiver Bandwidth: 83 kHz, Slice Thickness:1 mm, RARE Factor: 2.

#### 2.4.2 Data analysis

##### 2.4.2.1 Inversion efficiency

Inversion efficiency was calculated from the fc-FLASH images following reconstruction from the raw k-space data using an in-house script. Individual coil images were reconstructed for the control and label conditions, and α was calculated for each carotid and coil from the complex-valued reconstruction, according to:

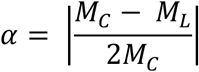

Where α is inversion efficiency and *M_C_* and *M_L_* are the extracted signals from the carotid arteries from the control and label conditions, respectively. Because two of the coil images exhibited low signal-to-noise ratio (SNR), inversion efficiency was calculated only from the two channels with the highest SNR. For each subject, the inversion efficiency was then defined as the mean value of the extracted signal from the right and left carotid arteries.

##### 2.4.2.2 T1 map

T1 maps of each subject were calculated by voxel-wise fitting of the saturation recovery equation to the signal obtained at various repetition times from the RARE-VTR sequence according to:

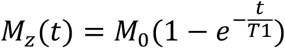

Where *M*_0_ is the equilibrium magnetization, *M_z_* is the longitudinal component of magnetization at each TR and T1 is the longitudinal relaxation time of tissue. All curve fitting was performed using the Levenberg-Marquardt algorithm using the SciPy Python package ^37^.

##### 2.4.2.3 CBF map

CBF (ml/100g tissue/min) were calculated on a voxel-by-voxel basis according to the single-compartment model of the general kinetic model for the pCASL signal ^38^:

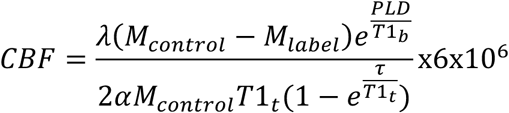

Where *λ* is the blood-tissue partition coefficient of water, *M_control_* and *M_label_* are the magnitude control and label images averaged across all repetitions, PLD is the post-labeling delay (ms), *T*1*_t_* (ms) is the longitudinal relaxation time of blood, calculated from the mean across all subjects, *T*1*_b_* (ms) is the longitudinal relaxation time of blood ^39^.

For analysis of CBF in different regions-of-interest (ROIs), pCASL-EPI images and Allen Institute Mouse Brain ^40,41^ annotations were registered into a common anatomical space. Control pCASL-EPI and MSME images were averaged across repetitions and echoes, respectively, and skull-stripped. All images underwent N4 bias field correction using the SimpleITK Python package ^42^ to reduce coil inhomogeneity. A group-level template was generated in ANTs ^43^, from the MSME slice corresponding to the pCASL-EPI acquisition position for each subject. The mean pCASL-EPI images and atlas template were aligned to each individual subject’s mean MSME image and then the group-level template by using a series of transformations: rigid-body, followed by affine transformations and finally non-linear SyN transformations using the ANTsPy (https://github.com/ANTsX/ANTsPy) Python package. The resulting transformations were applied to the calculated CBF maps and the atlas labels. From the transformed Allen Institute atlas labels, regional CBF values were extracted from a predefined set of ROIs present in the pCASL-EPI slice. CBF values below zero were thresholded to remove nonphysiological measurements. ROIs included the neocortex, SSC-BF, hippocampus, thalamus, and hypothalamus. Each ROI was subdivided by hemisphere, yielding combined bilateral and separate right and left hemisphere measurements for all regions.

##### 2.4.2.4 Cortical Thickness

Cortical thickness was calculated for each subject on the MSME slice corresponding to the pCASL-EPI acquisition position. Gray matter/white matter and gray matter/cerebrospinal fluid boundaries were manually delineated for each slice, and cortical thickness was computed using the LayNii toolbox^44^. Cortical thickness was separated by hemisphere to obtain combined bilateral, and separate right and left hemisphere measurements.

### 2.5 Immunofluorescence imaging

Mice were deeply anesthetized with pentobarbital (100 mg/kg i.p.). Perfusion was performed with 10-20 mL of phosphate buffered saline (PBS), followed by 4% paraformaldehyde (PFA) in PBS. The brains were removed and post-fixed in 4% PFA for 72 hours. The brains were cut into 50 μm coronal sections on a microtome (Leica SM2000R). Prior to staining free-floating sections were washed three times in 0.3% Triton X-100 in PBS. Sections were incubated in the blocking solution (protein block, abcam ab64226), followed by incubation with the primary antibody (anti-NeuN, 1:500 [abcam ab279297-1005], overnight at 4°C. After overnight incubation and washing, the sections were incubated in secondary antibody (1:1000, abcam ab150073-1001) for 2 hours at room temperature and followed by 1 hour incubation in DAPI (1μg/mL). The sections were mounted on the slides with fluoromount-G mounting media. Stained sections were imaged using a Zeiss Axioscan Z1 microscope with a 20x/0.8 NA plan apochromat objective.

### 2.6 Craniotomy and multiphoton imaging

Mice were anesthetized with isoflurane (5% for induction, 0.5-2% for maintenance) throughout the surgery. A custom built headplate was affixed with C&B Metabond dental cement (Parkell # S380) to the skull to immobilize the head during imaging. Body temperature was maintained at 37°C throughout surgery and imaging using a rectal probe feedback-regulated heatpad (Stoelting, Co.). A circular cranial window (∼ 3 mm diameter) was drilled over the somatosensory cortex (∼-1.5 mm AP, 3.0 mm ML vs. bregma). Texas red-dextran (70 kDa, 3 mg/ml, 150 μL) was injected into the retro-orbital sinus to enable visualization of cerebral vasculature. Upon conclusion of surgery, isoflurane anesthesia was replaced with 100mg/kg ketamine and 10 mg/kg xylazine. Artificial cerebrospinal fluid (aCSF; 124 mM NaCl, 3 mM KCl, 2 mM CaCl2, 2 mM MgCl2, 1.25 mM sodium phosphate buffer, 26 mM NaHCO3, and 4 mM glucose) was applied to the exposed cortex for the duration of the experiment.

Single-plane imaging experiments were performed in cortical layers II-III (∼250 µm depth) at 30 Hz for 6 min, with a FOV of approximately 556 x 556 µm. For a subset of experiments, 100 μM Barium chloride was perfused over the cortex for a minimum of 30-minutes to allow penetration. Images were acquired using Bruker Ultima 2P plus (Bruker, USA) with Nikon 16x plan apochromat 0.8 NA DIC VIS-IR water immersion objective, coupled with Coherent Discovery LX laser (Coherent, USA). GCaMP8 and Texas red were excited at 920 nm and emitted fluorescence was collected through bandpass filters (Chroma, ET525/50m-2p and ET595/50m2p) and GaAsP photomultiplier tubes (Hamamatsu Model H10770PB-40).

### 2.7 Widefield imaging

To express GCaMP7s pan-neuronally, mice received retro-orbital injections of PHP.eb-AAV-syn-jGCaMP7s-WPRE (Addgene, 104487-PHPeB) (1×10^13^ vg/ml, 7μl) at 8 weeks age-point. Around 11 weeks of age, mice were anesthetized with isoflurane (5% for induction, 0.5-2% for maintenance) for headplate implantation to stabilize the brain for imaging. The headplate was attached to the skull using C&B Metabond dental cement (Parkell # S380). For transcranial imaging, a thin layer of UV curing cyanoacrylate was applied on the surface of the skull. After surgery, the headplate was filled with silicone elastomer (Kwik-Sil, WPI) for protection during recovery and meloxicam (5 mg/kg, s.c.) was administered for post-operative pain management. Mice were allowed to recover for 3-4 days followed by three rounds of habituation to the imaging platform. Habituation sessions were performed as follows: day 1 – 10 minutes, day 2 – 20 minutes, day 3 – 30 minutes. Mice were secured under the objective lens and supported by a custom treadwheel during habituation and imaging. Transcranial imaging was performed for 3 consecutive days for 30 minutes each day in controls and 5xFAD mice to capture wide-field neuronal activity in the field of view (∼9 x 5 mm). Data were acquired at 30 Hz using an epi-fluorescent microscope (Bruker Ultima) with a Nikon 2x Plan Apo Lambda 0.10 NA objective and filter cube set ET-EGFP (FITC/Cy2) with filters ET470/40x and ET525/50m; dichroic T495lpxr. Behavioral data during imaging sessions were collected with infrared illumination using an Allied Vision Alvium 1800 camera with a Tamron 12VM412ASIR infrared lens and recorded with StreamPix software.

### 2.8 Imaging data analysis

#### 2.8.1 Analysis of multiphoton data

Images from multiphoton experiments were acquired at 30Hz. Recordings were condensed into 12-frame averaged files and further denoised using a 3×3 gaussian filter to remove Poisson noise. The images were then analyzed using an analysis pipeline reported previously with modifications^29^. First, the 12-frame averaged recordings were motion corrected using rigid registration to align each frame with the first. (*σ*_*n*_) Next, the standard deviation of each pixel’s intensity (*σ_n_*) was calculated over non-overlapping 30-frame spans; these successive values were compared by converting to imaging decibels (idB*_σ_*), defined below, to identify large changes in intensity.

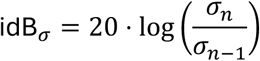

Pixels with |idB*_σ_*| ≤ 10 were removed. Remaining signals that take up more than 20 pixels in a maximum time-projection were classified as active sites. Active sites identified as non-capillaries (i.e., arterioles and venules) were manually erased based on large lumen size (>5-8μm) and directionality (vertical positioning through field). z-scores were calculated by assessing the fluctuations in intensity relative to quiescent background periods. The z-scores from each active site were extracted and plotted on a spatiotemporal (ST) map with a cutoff of 2.5. The length and duration of each active site event are represented in the ST map such that the x-axis represents the length of an event along the corresponding active site’s longest axis, and the y-axis represents the duration of elevated signal.

#### 2.8.2 Analysis of widefield imaging data

Widefield fluorescence data were processed to calculate the relative change in fluorescence from raw recordings. The baseline was defined as the average of the 20th percentile of pixel intensity over the recording duration, consistent with the established protocols^45^. To separate signal from noise, rank 50 Singular Value Decomposition (SVD) was applied to the spatial and temporal components^45^. The temporal components were detrended and bandpass filtered to remove physiological artifacts (e.g., heart rate) and slow signal drifts before reconstructing the denoised image stacks. Downstream analyses, including trace extraction, heat map visualization, and event detection, were performed on the reconstructed data.

##### Peak event detection

To characterize neuronal activity, the SVD-reconstructed 30 Hz signals were first down sampled to 0.2 Hz to smooth high-frequency noise and reveal macro-structural signal dynamics. A dynamic baseline was calculated as the 20th percentile of the smoothed signal. Neuronal events were defined as contiguous periods where the signal amplitude strictly exceeded this baseline. Following event identification, peak amplitude and timing were extracted from the SVD-reconstructed high-resolution (30 Hz) traces. The peak amplitude was determined as the maximal single-frame intensity within the identified time windows to ensure the reported magnitude was not dampened by down sampling. Parameters including event duration, peak time, and peak amplitude were aggregated for analysis.

##### Locomotion quantification

Locomotion was differentiated from stationary periods using an automated frame-by-frame motion detection pipeline. A Gaussian blur was applied to raw video frames to suppress high-frequency noise, followed by the generation of a difference map between consecutive frames. A binary threshold was applied to the difference map to isolate pixel changes, generating a time-series signal of motion intensity. To distinguish true locomotion from background noise and fine motor movements, a dynamic statistical threshold was calculated using Otsu’s method based on the bimodal distribution of pixel counts. Significant motion events were defined as intervals exceeding the threshold for a minimum of 1 second (30 frames). Distinct events occurring within a short interval (0.33 seconds/10 frames) were merged into single continuous epochs, and the timestamps and durations of these detected events were aggregated for analysis.

##### Atlas registration

A 2D reference atlas was generated from the Allen Brain Atlas CCFv3 (2017) by selecting the immediate volumetric nodes under the Isocortex (structure ID 315). These volumes were rendered to produce a dorsal view of the cortical surface, from which region boundaries were extracted. As the cranial window afforded a partial field of view (FOV), the atlas was manually registered to the experimental FOV using superficial vasculature and anatomical landmarks as reference points.

#### 2.8.3 Vessel segmentation and capillary density calculation

Vessel segmentation and capillary density measurement were performed with steps described below (**Supplement figure 1**).

##### Image Denoising

For capillary density analysis, three-dimensional (3D) images were denoised with SUPPORT, a self-supervised deep learning method which removes Poisson and Gaussian noise in static volumetric structural imaging^46^. Training and inference were conducted independently for each volume. The denoising networks were trained using Pytorch 2.5.1 on an NVIDIA A100 Tensor Core GPU.

The same hyperparameters were used to denoise all volumes: blind plane disabled, 3 x 3 blind spot size, 128 batch size, and a learning rate of 5×10^-6^. Models were saved after training for 150 epochs. For training, 3D patches of size 128(*x*) x 128(*y*) x 61(*z*) and interval 64(*x*) x 64(*y*) x 1(*z*) were extracted from the input image. To preserve the input shape, we added reflective padding of 30 to the first and last slice prior to inference. For inference, the patch interval was reduced to 11(*x*) x 11(*y*) x 1(*z*) to mitigate seam artifacts.

##### Vessel Segmentation

Next, we segmented the vessels in each of the denoised images using VesselFM, a foundation model pretrained for universal 3D blood vessel segmentation^47^. To isolate capillaries from large vessels, we removed objects thicker than 8µm (in diameter) and longer than 30µm. Subsequently, we removed using a 144-voxel small-particle filter using scikit-image’s (v0.26) morphological algorithm^48^. This value was determined by computing the volume corresponding to the smallest range of capillary lengths and radii required to visually distinguish segmented capillaries from noise.

##### Capillary Density Measurements

Capillary density (*⍴*_c*l*_) is defined as the sum of all capillary lengths *l*_*n*_, divided by the imaged volume *V*.

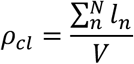

Where *N* is the total number of capillaries identified in the vessel segmentation step. Measuring the length of capillaries starts with computing the skeleton of the capillary mask using scikit-image’s skeletonize function^48^. Another Python package, Skan (v0.13), then computes capillary lengths by summing the Euclidean distances (scaled by voxel resolution) between adjacent pixels along each skeletonized branch^49^.

**Supplement figure 1:**
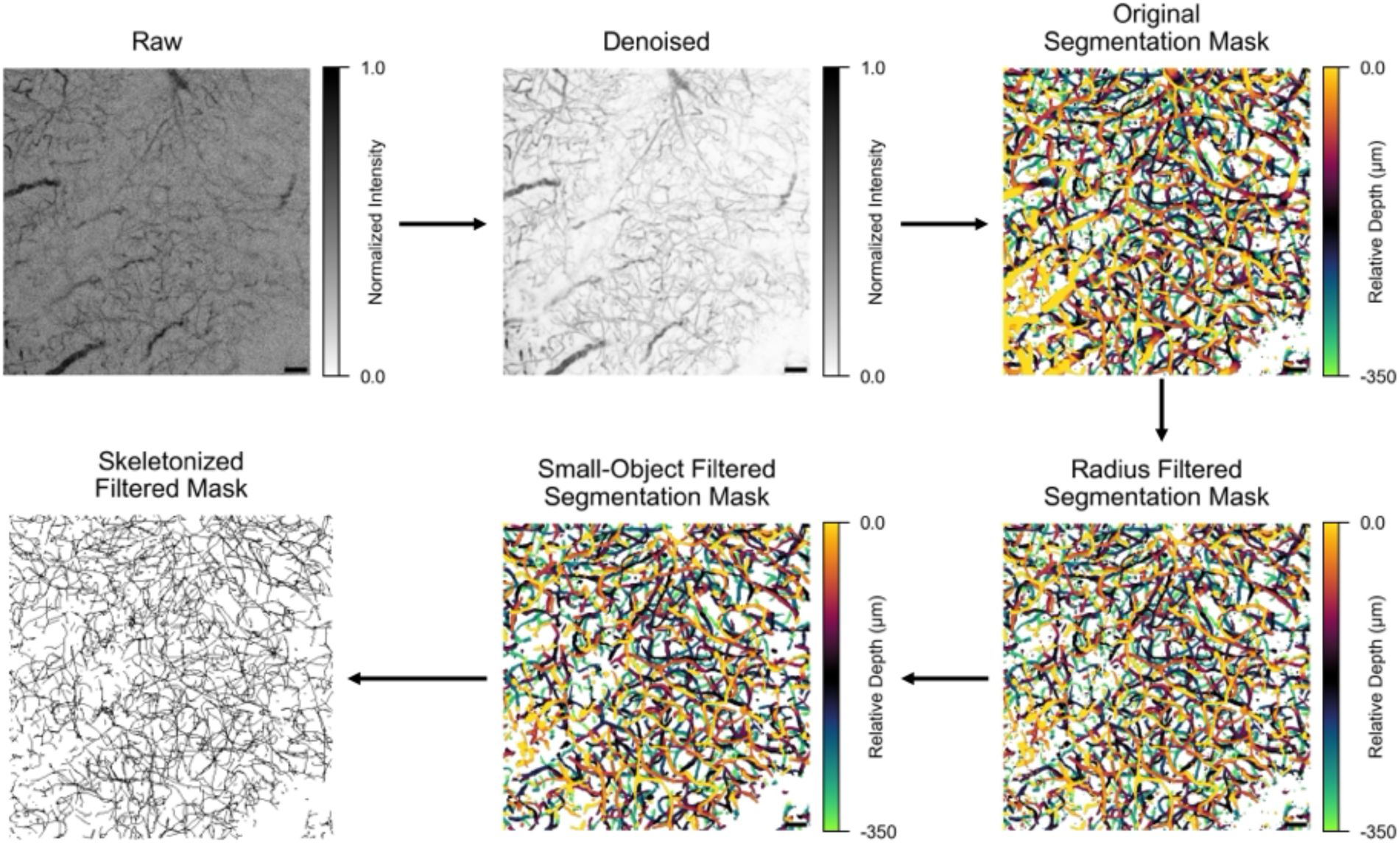
Representative maximum z-projections illustrating processing steps used for capillary density measurements shown sequentially clockwise. The raw data is first denoised and segmented. Capillaries are isolated by removing objects with diameter >8µm and volume <144 voxels. The skeleton of the filtered capillary mask is used to calculate their length.

#### 2.8.4 Neuronal density measurements

Cortical neuronal nuclei were quantified using Fiji (ImageJ) v2.14 on 8-bit images. Images were preprocessed with the background subtraction and Otsu intensity threshold to create a binary mask. Nuclei segmentation was performed with the watershed segmentation approach, and the segmented nuclei were counted with analyze particle function with criteria for circularity: 0.2-1.0 and size: 50-300 μm^2^. The number of neuronal nuclei within the dorsal neocortex was divided by the area of the region to obtain the neuronal density.

### 2.9 Electrophysiology

To evaluate the deficits in electro-calcium coupling in capillary endothelial cells, ex-vivo patch clamp electrophysiology was performed using conventional whole-cell voltage clamp method at room temperature (∼22°C). Currents were amplified using Axopatch 200B (Molecular Devices) and the signal was low-pass filtered at 1 kHz and digitized at 10kHz using Digidata 1550B (Molecular Device). Upon breaking into cell, whole cell capacitance (C_m_) and series resistance (R_s_) was measured and compensated using the cancellation circuitry (C_mem_ 8.55 ± 0.4 pF). Patch with R_s_ greater than 15MΩ was excluded from analysis. Patch pipette were fabricated from borosilicate glass (OD/ID = 1.5/1.17 mm, Sutter instruments) using P-1000 Horizontal puller (Sutter instruments) and fire-polished using MF-2 microforge (Narishige) to reach pipette resistance of ∼2-4MΩ, and filled with internal solution consisting of 10 NaOH, 11.4 KOH, 128.6 KCl, 1.1 MgCl_2_, 2.2 CaCl_2_, 5 EGTA, and 10 HEPES (pH7.2). For K_ir_2.1 experiments, bath solution consisted of 80 NaCl, 60 KCl, 1 MgCl_2_, 2 CaCl_2_, 4 D-glucose, and 10 HEPES (60 K^+^, pH 7.4). For TRPV4 experiments, bath consisted of 134 NaCl, 6 KCl, 1 MgCl_2_, 2 CaCl_2_, 4 D-glucose, and 10 HEPES (6 K^+^, pH 7.4). Pharmacological inhibitors (100 µM BaCl_2_ for K_ir_2.1 and 1 µM Ruthenium Red for TRPV channels) and activators (1 µM GSK1016790A for TRPV4, GSK101 hereafter) were made fresh on the day of the experiment by diluting 1000x stock in respective bath solutions.

#### 2.9.1 Cell isolation protocol

On the day of the experiment, we performed ex-vivo isolation following previously established protocol ^26,27,50^. Animals were sacrificed and brains were extracted into ice-cold aCSF. A small coronal slice (∼160 µm) of cortex (∼2 mm posterior to the bregma) was taken and homogenized in ice-cold aCSF using Dounce homogenizer. The cell suspension was filtered through 62 µm nylon filter (Component Supply) and the remaining cell fragments were enzymatically digested using 0.5 mg/ml of neutral protease and elastase (Worthington) in dissociation buffer (55 NaCl, 80 Na-Glutamate, 5.6 KCl, 2 MgCl_2_, 4 D-glucose, and 10 HEPES pH 7.4) for 24 min at 37°C. The tissue was then further digested using 0.5 mg/ml of collagenase type 1 (Worthington Biochemical) and incubated for additional 2 min at the same temperature. The cell suspension was filtered using the same filter and washed using dissociation buffer. The remaining fragments were diluted in dissociation buffer and triturated using fire-polished glass Pasteur pipette (∼4-6x). The cells were kept ice-cold and used within 6 hours of isolation. The cell suspension was added dropwise (∼10-15 drops) to a 1 ml chamber filled dissociation buffer and the cells were allowed to adhere to the chamber for ∼40 min prior to recordings. The bath solution was exchanged with respective bath as described above (60 K^+^ or 6 K^+^).

#### 2.9.2 Data acquisition and analysis

The currents were evoked using a ramp protocol with a ramp velocity of 1 V/s. For both K_ir_2.1 and TRPV4, the ramp duration was 200 ms and held at initial and final step voltage for 25 ms prior termination of the sweep. For both protocols, holding potential was set at −50 mV. For K_ir_2.1 single sweep with ramp from −150 to 50 mV was used. To isolated K_ir_2.1 current, the recording in 60 K^+^ bath was subtracted by the recording in 60 K^+^ bath supplemented with 100 µM BaCl_2_. To evaluate the steady-state current density (Current Density_Kir2.1_), the average of 5 ms (51 data points at 10 kHz) at step voltage of −150 mV was normalized to the cell capacitance (C_m_) measured for each cell. For TRPV4 ramp was from −100 mV to 100 mV and sweep was repeated every 10 seconds for 10 minutes in presence of GSK101 to allow sufficient time for the current to saturate. To evaluate the steady-state current density of TRPV4, GSK101 saturated current at +100 mV step was averaged for 5 ms and normalized to the cell capacitance as above.

### 2.10 Statistics

Statistical testing was performed using GraphPad Prism 10 software. Data are expressed as means ± standard error of the mean (SEM), and a p-value < 0.05 was considered significant. Data were tested for normality using a Shapiro–Wilk test prior to application of the appropriate parametric or nonparametric statistical test. Stars denote significant differences. Statistical tests are noted in figure legends, where ‘ns’ represents not significant differences and ‘n’ refers to the number of animals used, unless otherwise stated. All t-tests were two-sided. Sample sizes were estimated based on the power analysis and/or similar experiments previously performed by the lab. Data collection was not performed blinded to the conditions of the experiments. Littermates were randomly assigned to experimental groups; no further randomization was performed.

## 3. RESULTS

### 3-month-old 5xFAD mice do not have cognitive impairment

5xFAD mice are reported to have irreversible neurodegeneration and cognitive decline around 4-9 months of age ^51^. We subjected young adult (3-month-old) 5xFAD mice and age-matched controls to a battery of behavioral tests to verify that these mice were presymptomatic or in a mild cognitive impairment (MCI)-like stage, suitable for assessing early neurovascular coupling (NVC) deficits.

First, we evaluated spatial learning and memory with Y-maze and NOR. The design of these two tests allowed us to identify deficits in short term recall (Y-maze) and long-term memory consolidation (NOR). In Y-maze test, the mice were placed in Arm A and allowed to explore three different arms (Arms A, B, and C) to evaluate their ability to spontaneously alter (any combination of three consecutive visit to an arm without overlap) between these arms (% alteration) as a measure of their short-term recall (**Figure 1A**). 5xFAD mice and their age-matched controls exhibited comparable spontaneous alteration between arms (**Figure 1B-C**) indicating intact short-term recall. Next, we evaluated long-term memory consolidation with NOR. In NOR test, the mice were exposed to two same objects for 10 minutes to become familiar with these objects and consolidate this memory for 24 hours. After 24 hours, one of these familiar objects was replaced with a novel object and mice were exposed to a novel and a familiar object for 10 minutes. The long-term memory consolidation was evaluated based on their ability to identify the novel object and explore more time with this object (**Figure 1D**). Both control and 5xFAD mice were able to distinguish novel object from the familiar object and spent more time with the novel object as presented with the discrimination index (**Figure 1E**), suggesting no change in the long-term memory consolidation. Lastly, we assessed anxiety-like state and motor functions with the open field test. Where each group was given 10 minutes to explore an open-rectangular box (dimensions: 46 x 46 x 50 cm box). Time spent in the center area (23 x 23 cm) of the box (**Figure 1F**) and total distance traveled were used to measure the anxiety-like behavior and motor function. Both 5xFAD mice and their respective age-matched controls spent similar time in the center and traveled equal distance suggesting absence of anxiety-like behavioral or motor deficits (**Figure 1G and H**). Altogether, these data confirmed that 3-month-old 5xFAD mice do not present with the short-term recall, long-term memory consolidation, and anxiety-like behavior deficits.

**Figure 1.**
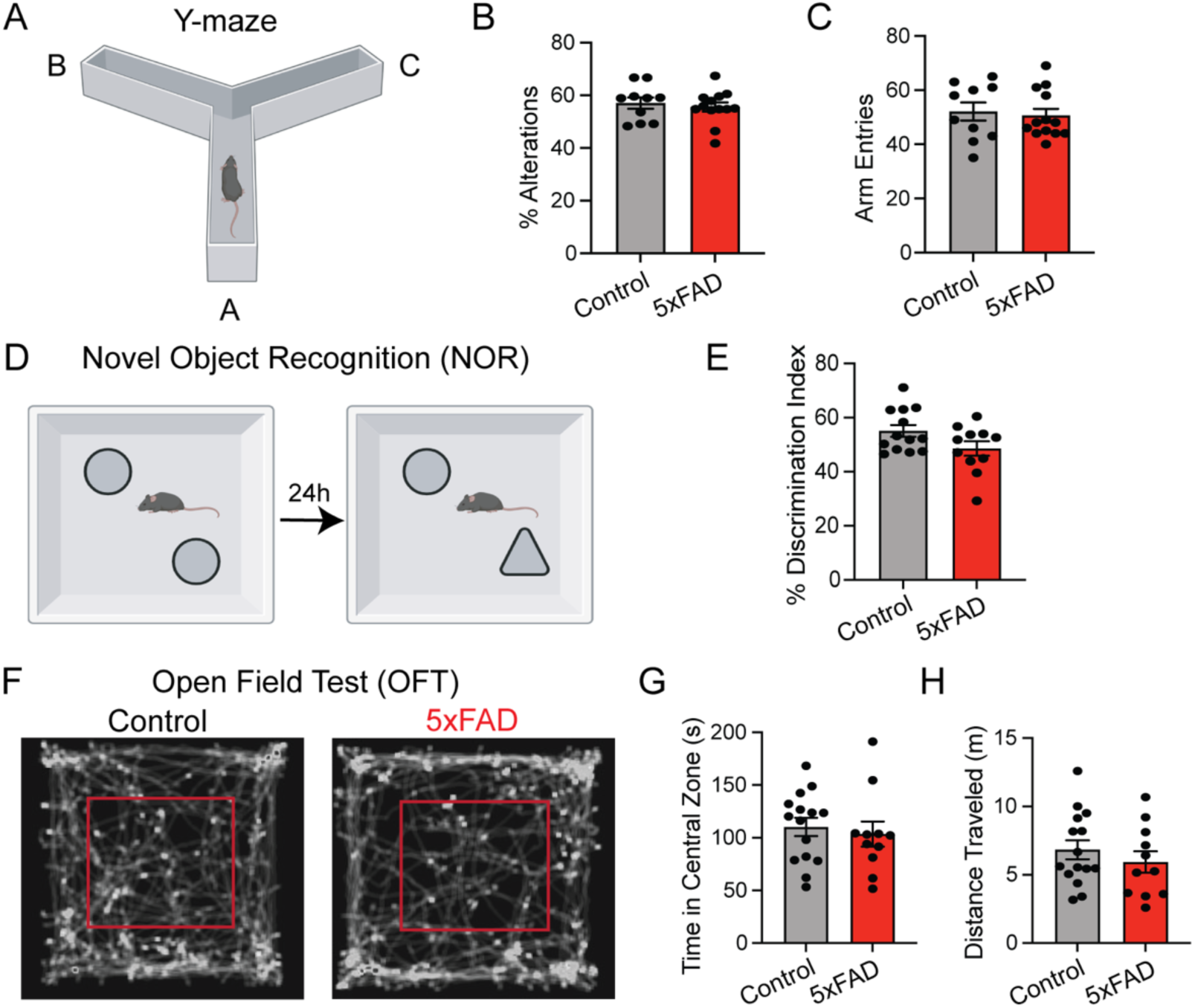
Young 5xFAD mice do not present with cognitive impairment. **(A)** Schematic representing Y-maze set-up, **(B-C)** Y-maze test showed 5xFAD mice have comparable **(B)** spontaneous alterations (control: 57.0 ± 2.15% vs. 5xFAD: 55.6 ± 1.75%, ns, n=10-13, unpaired t-test) and **(C)** arm entries (control: 52 ± 3 vs. 5xFAD: 50 ± 2 entries, ns, n=10-13, unpaired t-test) as controls. **(D)** Schematic representation of novel object recognition (NOR). **(E)** No significant difference was observed in the discrimination index between controls and 5xFAD animals (control: 55.07 ± 2.2% vs. 5xFAD: 48.54 ± 2.66%, ns, n=11-13, unpaired t-test). **(F)** Representative locomotion traces of control (left) and 5xFAD (right) mice in the open field test. Red boxes indicate center of the open field. **(G-H)** Time in central zone (control: 110.3 ± 8.36 vs. 5xFAD: 103.5 ± 11.87 sec, ns, n=10-15, unpaired t-test) and distance traveled (control: 6.82 ± 0.69 vs. 5xFAD: 5.93 ± 0.80 m, ns, n=10-15, unpaired t-test).

### Functional hyperemia, but not the resting-state perfusion is impaired in 3-month-old 5xFAD mice

With our goal to identify NVC deficits during early presymptomatic phase, we performed cerebral hemodynamic measurements in 3-month-old 5xFAD mice and age matched controls. We hypothesized that these 5xFAD mice have a significant reduction in the functional hyperemia. To test this hypothesis, we measured whisker stimulation (WS)-induced increase in cerebral blood volume (CBV) in the barrel field of somatosensory cortex (SSC-BF). We used thinned-skull anesthetized mice and fUS to measure changes in CBV with a WS protocol that consisted of two 30-sec long stimulations separated by a 60-sec baseline. During the WS period, air puffs (20 psi at 20 Hz) were given and activation maps were computed using Pearson-correlation coefficient. This WS-paradigm led to an increase in CBV in the contralateral cortex, while ipsilateral cortex served as a built-in experimental control (**Figure 2A**). The time course of normalized CBV change (rCBV) shows robust increase in steady-state CBV upon WS in the control mice (rCBV: 16 ± 0.55 %), whereas 5xFAD mice showed a significant reduction (rCBV: 11± 1.2 %) (**Figure 2B-C**). Further temporal analysis showed that WS-induced increase in CBV follows a bimodal response, consisting of an initial fast increase in CBV (P_fast_) followed by a prolonged response reaching to the steady state (P_slow_) (**Figure 2D**). The bimodal response of functional hyperemia has been described in previous reports with awake behaving and anesthetized rodent models^30,52–54^. However, we are the first to characterize vascular components involved in each phase and extend this to 5xFAD mice. Notably, both control and 5xFAD mice showed the bimodal response, albeit with key differences in the two phases. To compare these differences, we fit single-exponential functions to quantify the kinetics of the fast (P_fast_), slow (P_slow_), and decay phase (P_dec_). Our image acquisition rate (2.5 Hz) was relatively slower compared to the rapid kinetics of the fast phase, P_fast_ fitting had an insufficient temporal resolution for analysis. To overcome this limitation, we quantified overall duration of the fast phase. The rise time constants for the slow phase (τ_slow_; Control: 3.1± 0.98 *vs.* 5xFAD: 2.3 ± 0.56 sec) and decay time constants (τ_dec_; Control: 1.6 ± 0.42 *vs.* 5xFAD: 2.1 ± 0.45 sec) did not significantly differ between control and 5xFAD mice (**Figure 2E**). However, duration of the fast phase was slightly longer in 5xFAD mice (**Figure 2F**). Moreover, the fast-to-slow phase transition was markedly impaired in 5xFAD mice (**Figure 2D, bottom**). This transition impairment was quantified using amplitude drop (A_drop_) and difference in the transition time (ΔT_drop_). Both A_drop_ (Control: −2.2 ± 0.51 vs. 5xFAD: −7.4 ± 1.6%) and ΔT_drop_ (Control: 0.29 ± 0.14 *vs.* 5xFAD: 1.2 ± 0.31 sec) were significantly larger in 5xFAD mice compared to their age-matched controls (**Figure 2G and 2H**). These differences suggest that functional hyperemia deficits in 3-month-old 5xFAD mice result from impaired fast-to-slow phase transition.

**Figure 2.**
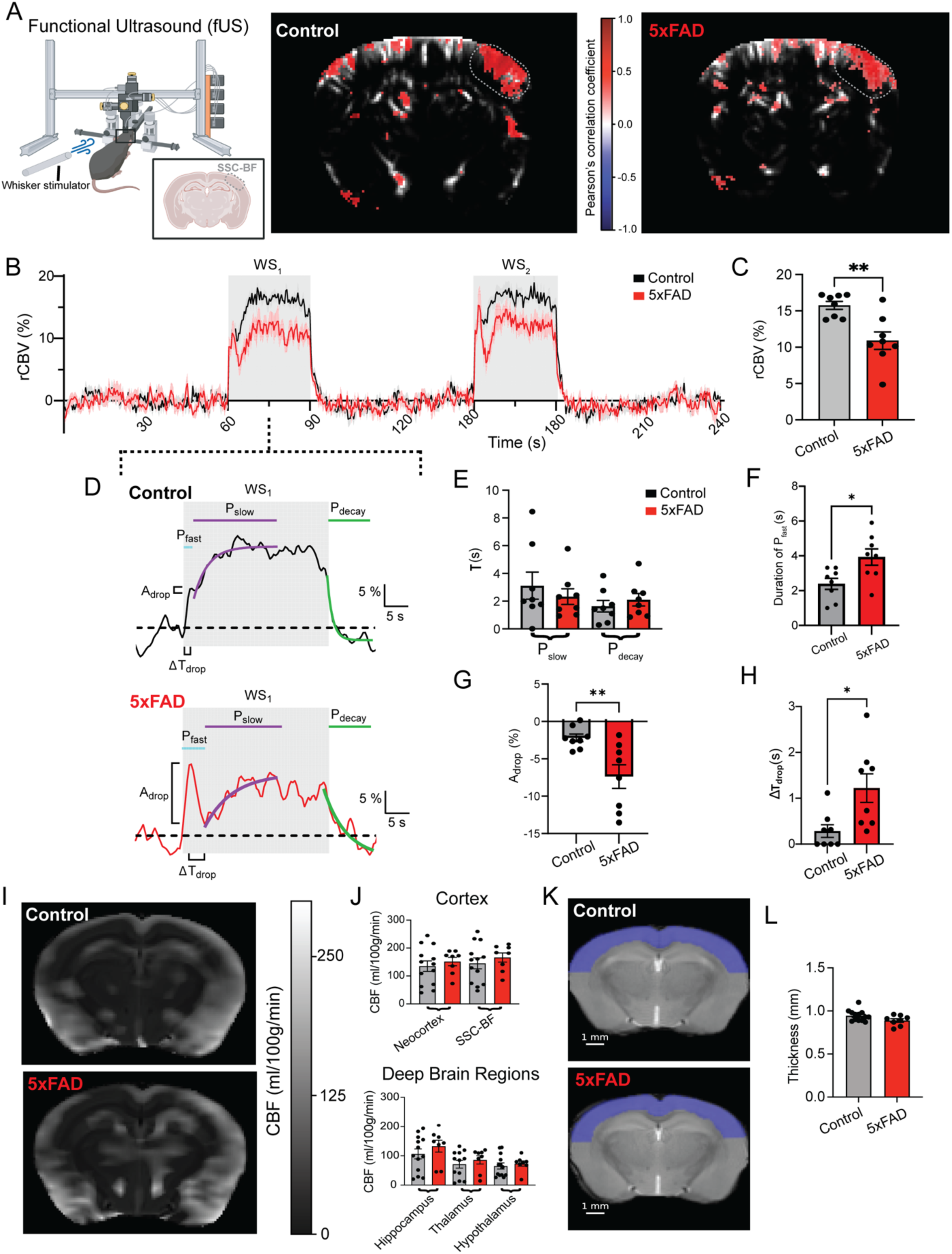
5xFAD mice exhibit deficits in functional hyperemia but not in the resting-state blood flow. **(A)** Schematic representing Functional ultrasound (fUS) imaging and whisker stimulation set-up along with the coronal slice of interest encompassing somatosensory cortex, Barrel field (grey-dotted oval) (left). Activation map generated with Pearson’s correlation coefficient in response to left-side whisker stimulation, with grey-dotted oval denoting contralateral activation, for control (middle) and 5xFAD (right) shown. **(B)** Time course of normalized rCBV change in control (black) and 5xFAD (red), the duration of recording is 240s with two 30s stimulation period (grey shaded area). **(C)** Comparison of steady-state change in rCBV at first whisker stimulus period (WS_1_) between control and 5xFAD. The 5xFAD mice show reduction in steady-state rCBV change at 3-month-old. (Control: 16 ± 0.55 vs. 5xFAD: 11 ± 1.2 %, **p<0.01, unpaired t-test, n=8). **(D)** Representative trace of control (top) and 5xFAD (bottom) showing WS_1_ period, note the three phases, fast rise kinetics (P_fast_, light blue), slow rise kinetics (P_slow_, purple), and decay kinetics (P_decay_, green). **(E)** Comparison of the time constant (τ) for control and 5xFAD from exponential fit of the slow (Control: 3.1 ± 0.98 vs. 5xFAD: 2.3 ± 0.56 sec, ns, unpaired t-test, n=8) and decay kinetics (Control: 1.6 ± 0.42 vs. 5xFAD: 2.1 ± 0.45 sec, ns, unpaired t-test, n=8), respectively. **(F)** Comparison of the fast phase duration between control and 5xFAD mice (Control: 2.4 ± 0.32 vs 5xFAD: 3.9 ± 0.46 sec, * p<0.05, n=8). **(G and H)** Comparison of two transition impairment parameters: **(G)** amplitude drop (A_drop,_ Control: −2.2 ± 0.51 vs. 5xFAD: −7.4 ± 1.6 %, ** p<0.01, unpaired t-test, n=8) and **(H)** difference in transition time between fast and slow phase (ΔT_drop_, Control: 0.29 ± 0.14 vs. 5xFAD:1.2 ± 0.31 sec, *p<0.05, unpaired t-test, n=8). **(I)** Representative CBF maps for control (top) and 5xFAD (bottom) overlaid on top of anatomical images obtained from MRI. **(J)** Comparison of average CBF of cortex (top; neocortex; control: 135 ± 19 vs. 5xFAD: 152 ± 15 ml/100g/min, ns; SSC-BF; control: 145 ± 19 vs. 5xFAD: 166 ± 17 ml/100g/min, ns, unpaired t-test, n=7-13) and deep brain regions (bottom; hippocampus; control: 107 ± 17 vs. 5xFAD: 133 ± 20 ml/100g/min, ns; thalamus; control: 71 ± 13 vs. 5xFAD: 86 ± 14 ml/100g/min, ns; hypothalamus; control: 65 ± 10 vs. 5xFAD: 74 ± 9.1 ml/100g/min, ns, unpaired t-test, n=7-13). **(K)** Representative anatomical image of control (top) and 5xFAD (bottom) coronal section with blue shaded area indicating the mask of area used for cortical thickness measurements. **(L)** Comparison of cortical thickness between control and 5xFAD (control: 0.94 ± 0.018 vs. 5xFAD: 0.89 ± 0.021 mm, ns, unpaired t-test, n=7-13).

Given that the resting cerebral perfusion can influence the functional hyperemia, we next measured resting cortical perfusion using pseudo-continuous arterial spin labeling-fMRI (pCASL-fMRI). pCASL-fMRI involves magnetically labeling inflowing arterial blood upstream of the imaging slice with RF-pulses, followed by acquisition of control and label images after a post labeling delay. The difference between the control and label image was used in conjunction with cerebral blood flow (CBF) equation (see methods) to quantify the absolute CBF. With this non-invasive approach, we measured cortical (neocortex and SSC-BF) and deep brain regions (hippocampus, hypothalamus and thalamus) cerebral perfusion (**Figure 2I**). We selected these regions as cortico-hippocampal pathways are critical for the memory consolidations and are shown to be impaired in AD ^55^. We used an anatomical atlas from the Allen Brain Institute ^40,41^ to register the imaging data, and atlas-defined regions were used to extract voxel-wise CBF measurements. In our measurements, we did not find any significant differences in resting-state cerebral perfusion between control and 5xFAD mice for both cortical (CBF_neocortex_: 135 vs. 152 and CBF_SSC-BF_: 145 vs. 166 ml/100g/min for control and 5xFAD respectively) and deep brain regions (CBF_hippocampus_: 107 vs. 133, CBF_thalamus_: 71 vs. 86, and CBF_hypothalamus_: 65 vs. 74 ml/100g/min for control and 5xFAD respectively) (**Figure 2J, top and bottom**) showing that the resting perfusion is comparable between control and 5xFAD mice. Moreover, reduction in the cortical thickness can also alter the cerebral perfusion. Thus, we measured cortical thickness using a high-resolution (in-plane resolution of 100 µm) coronal slice of the brain generated with MSME sequence (**Figure 2K**). Multi-point analysis across the length of neocortex showed that cortical thickness is similar between control and 5xFAD mice (**Figure 2L**). Collectively, these results demonstrate that 3-month-old 5xFAD mice exhibit a significant reduction in functional hyperemia, while the resting-state cerebral perfusion remains unchanged.

### Reduced functional hyperemia is linked with impaired capillary endothelial cell but not with neuronal activity

Functional hyperemia is driven by NVC that involves coordinated signaling between neural cells and blood vessels. We and others have previously described multiple NVC mechanisms that involve close communication between neural cells and capillary endothelial cells (cECs) ^11,26,30–32,53,56^. Out of these NVC mechanisms, cECs Ca^2+^ signals have relatively slow spatiotemporal kinetics for maintaining CBF during prolonged stimulation ^29^. Since, we observed a significant impairment in fast-to-slow phase transition functional hyperemia in response to whisker stimulation in 5xFAD mice (**Figure 2A-B**), we hypothesized that reduction in cECs Ca^2+^ activity contributes to impaired functional hyperemia in 5xFAD mice. We measured spontaneous cECs Ca^2+^ activity with genetic encoded Ca^2+^ indicator mice; *Cdh5*-GCaMP8 cross-bred with 5xFAD mice. We equipped a small circular cranial window over the somatosensory cortex of these mice and measured cortical cECs Ca^2+^ activity (layer II-III of the cortex) in a 556 x 556 mm field of view using multiphoton microscopy under ketamine and xylazine anesthesia (**Figure 3A**). Our measurements showed ∼40% reduction in the overall Ca^2+^ activity in the 5xFAD mice compared to their age-matched controls (control: 7303 ± 760 vs. 5xFAD: 2851 ± 635 zscr.μm.s.min^-1^) (**Figure 3B-D, Supplement movie S1-S2)**. Moreover, the total number of Ca^2+^ events generated in 5xFAD mice were significantly lower than their age-matched controls (**Figure 3E**). Stratification of these events in 3 different categories (proto, unitary and compound) based on the event duration^29^ showed that unitary and compound events were significantly reduced in 5xFAD mice, but proto events were similar between control and 5xFAD mice (**Supplement figure 2A-B**). These findings showed that functional hyperemia impairment in 5xFAD mice is linked with reduction in cECs Ca^2+^ activity.

**Figure 3.**
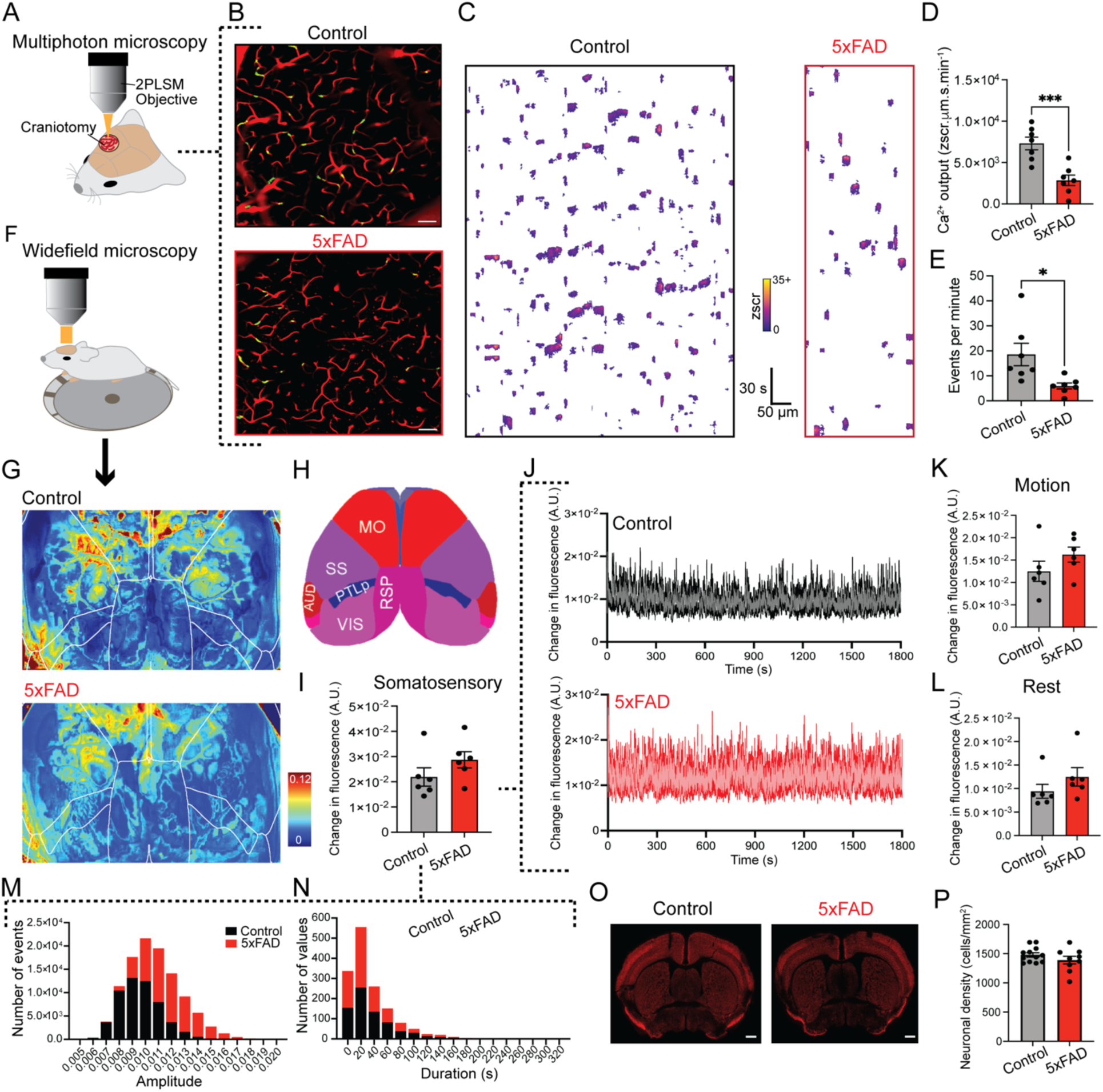
Capillary ECs Ca^2+^ activity are decreased, but neuronal Ca^2+^ dynamics and structure are unchanged in 5xFAD mice. **(A)** *In-vivo* imaging strategy with multiphoton microscopy. **(B-C)** Representative images where green and red indicate Ca^2+^ signals and plasma signals in the capillaries; respectively (FOV: 556 x 556 μm, scale bar: 50 μm) **(B)** and spatiotemporal (ST) maps **(C)** from control (left) and 5xFAD (right) mice. Scale bars apply to both maps. **(D)** cECs Ca^2+^ activity (control: 7304 ± 760 vs. 5xFAD: 2852 ± 635.4 zscr·μm·s·min^-1^, n = 7 experiments, 7 mice; ***p<0.001, unpaired t-test) and **(E)** total events per minute (control: 18.56 ± 4.48 vs. 5xFAD: 5.95 ± 4.48 n = 7 experiments, 7 mice; *p<0.05, unpaired t-test) in controls and 5xFAD mice. (F) *In-vivo* imaging strategy with wide-field microscopy. **(G)** Representative wide-field heat maps from control and 5xFAD mice aligned with **(H)** Allen Brain Altas (V3). **(I)** Total fluorescence changes (control: 0.022 ± 0.004 vs. 5xFAD: 0.028 ± 0.003 A.U., ns, n=6, Mann-Whitney test), **(J)** Time-traces showing change in fluorescence over time and changes in fluorescence stratified into **(K)** motion (control: 0.013 ± 0.002 vs. 5xFAD: 0.016 ± 0.002 A.U., n=6, ns) and **(L)** rest phase in somatosensory neurons of control and 5xFAD mice (control: 0.0094 ± 0.001 vs. 5xFAD 0.012 ± 0.002 A.U., ns, n=6). **(M-N)** Frequency histograms representing amplitude **(M)** and duration **(N)** of events in control and 5xFAD mice (n=6). **(O-P)** Representative images showing neuronal staining in coronal brain slices (scale bar= 500 mm) **(O)** and neuronal density **(P)** in control and 5xFAD mice (n=3-4 animals and 3 sections per animal, unpaired t-test).

Given that neurons initiate NVC and in turn, functional hyperemia, we next investigated neuronal contributions in reduced cECs Ca^2+^ activity and functional hyperemia in 5xFAD mice. To measure neuronal Ca^2+^ activity, we genetically encoded Ca^2+^ indicator jGCaMP7s in all neurons using PHP.eb AAV syn-jGCaMP7s. We performed transcranial imaging by rendering the bone semi-transparent with a layer of cyanoacrylate glue and imaged cortical GCaMP7s fluorescence with an epifluorescence microscope in awake head fixed mice freely running on a custom treadwheel (**Figure 3F**). We used Allen Mouse Common Coordinate Framework v3 (CCF) to align our imaging FOV and measured fluorescent intensity changes over time for multiple cortical regions (SS: somatosensory, PTLp: posterior parietal, MO: somatomotor and RSP: retrosplenial cortex) (**Figure 3G-H**). Since, functional hyperemia and vascular Ca^2+^ activity were measured in the somatosensory cortex, we centered FOV over the somatosensory cortex and also included other cortical regions in the analysis from the FOV. Total neuronal Ca^2+^ intensity in the somatosensory cortex did not differ between control and 5xFAD mice (**Figure 3I-L, Supplement movie S3-S4**) and other cortical regions (**Supplement figure 3A-F**). However, fluorescent intensity over time showed a subtle increase in high amplitude spikes in the somatosensory cortex of 5xFAD mice compared to the controls (**Figure 3J**). Frequency histograms of event amplitude and duration revealed that 5xFAD mice exhibited a higher proportion of large-amplitude Ca^2+^ events in the somatosensory cortex, whereas control mice displayed more frequent low-amplitude events. (**Figure 3M**), while duration of these events remained comparable between control and 5xFAD mice (**Figure 3N**). To evaluate the effect of locomotion on overall neuronal Ca^2+^ activity, we further stratified these data into motion and stationary phase. Total neuronal Ca^2+^ activity for motion and stationary phase was similar in the controls and 5xFAD mice (**Figure 3K-L, Supplement figure 3)**. Overall, these data indicate that reduced capillary vascular response—rather than changes in the neuronal activity—attenuates functional hyperemia in early age of 5xFAD mice.

**Supplement figure 2.**
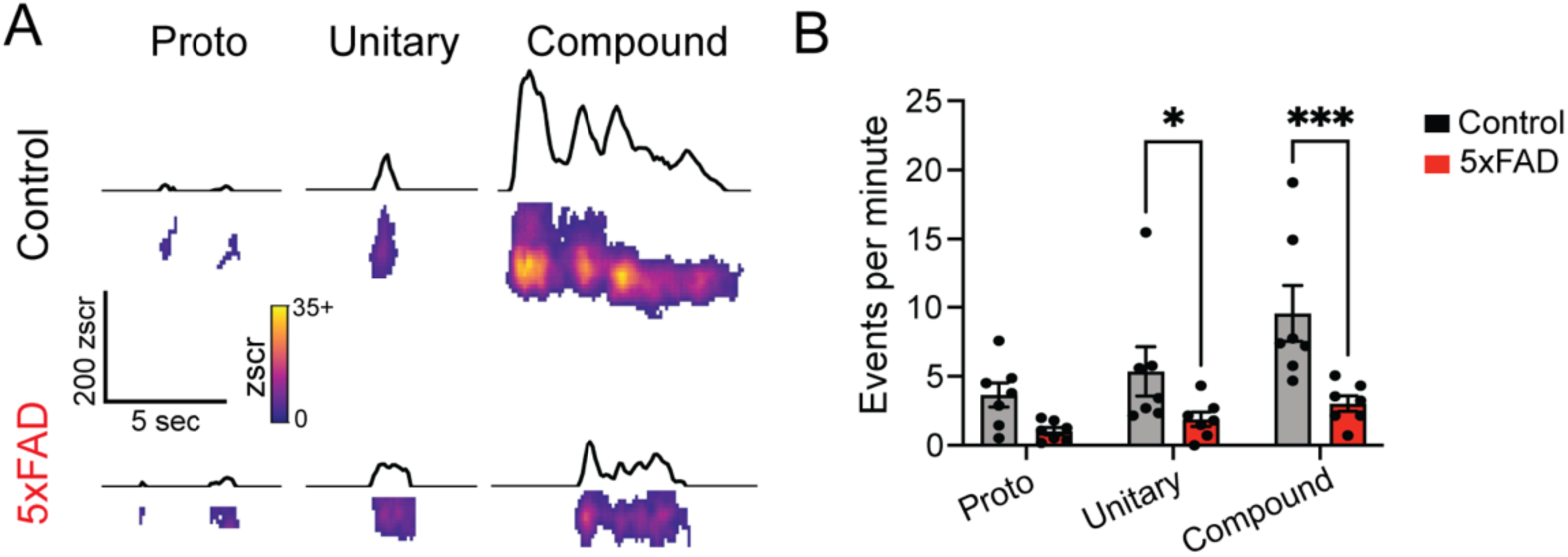
Characterization of cECs Ca^2+^ events in control and 5xFAD mice. **(A)** Representative traces of cECs Ca^2+^ events ranging from proto events to unitary and multicomponent compound events from controls (top) and 5xFAD mice (bottom). **(B)** Comparison of events by class in control and 5xFAD mice (proto, control: 3.7 ± 0.9 vs. 5xFAD 1.0 ± 0.3, ns; unitary control: 5.4 ± 1.8 vs 5xFAD: 1.9 ± 0.5, *p<0.05; compound, control 9.6 ± 2.0 vs 5xFAD: 3.0 ± 0.6 events·min^-1^, ***p<0.001; two-way ANOVA and Tukey’s multiple comparisons test, n=7).

**Supplement figure 3.**
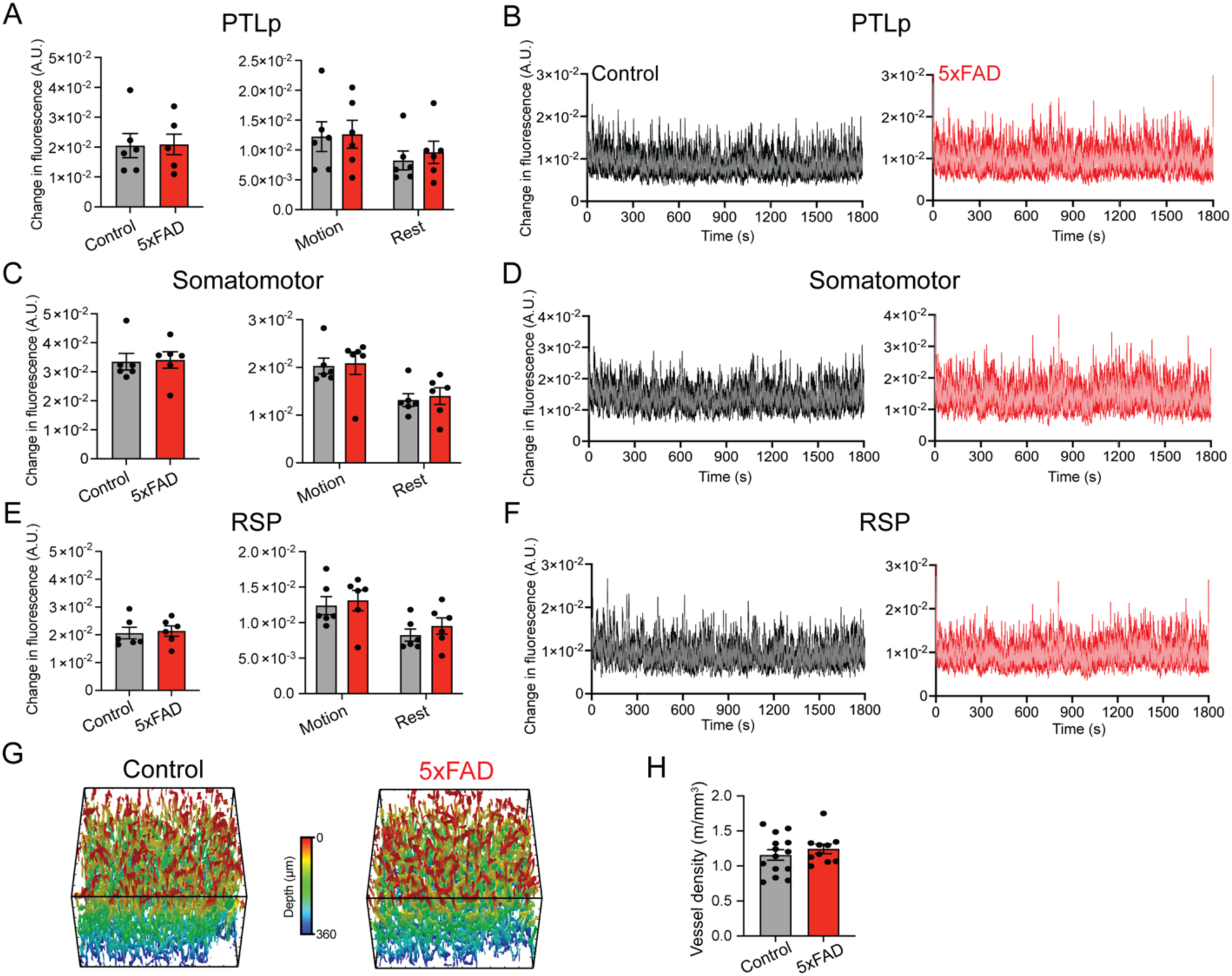
Neuronal Ca^2+^ activity and capillary density are unchanged in 5xFAD mice. **(A-F)** Neuronal Ca^2+^ presented as total activity, stratified into motion and stationary phase and time trace from control and 5xFAD mice for **(A-B)** PTLp: posterior parietal (control total: 0.020 ± 0.004 vs. 5xFAD total: 0.02 ± 0.003; control motion: 0.012 ± 0.008 vs. 5xFAD motion: 0.013 ± 0.002; control rest: 0.008 ± 0.002 vs. 5xFAD rest: 0.010 ± 0.002 A.U., ns, n=6, unpaired t-test), **(C-D)** MO: somatomotor (control total: 0.033 ± 0.003 vs. 5xFAD total: 0.034 ± 0.003, control motion: 0.020 ± 0.002 vs. 5xFAD motion: 0.021 ± 0.002, control rest: 0.013 ± 0.001 vs. 5xFAD rest: 0.014 ± 0.002 A.U., ns, n=6, unpaired t-test) and **(E-F)** RSP: retrosplenial cortex (control total: 0.021 ± 0.002 vs. 5xFAD total: 0.021 ± 0.002, control motion: 0.012 ± 0.001 vs. 5xFAD motion: 0.013 ± 0.001, control rest: 0.008 ± 0.001 vs. 5xFAD rest: 0.010 ± 0.001 A.U., n=6, ns, unpaired t-test). **(G-H)** Representative 3D-maps of capillary density in **(G)** control (left) and 5xFAD mice (right) and **(H)** comparison of capillary density between control and 5xFAD mice (control: 1.15 ± 0.07 vs. 5xFAD: 1.24 ± 0.07 m/mm^3^, ns, unpaired t-test, n=6-7 with two z-stacks per mouse).

### Cortical capillaries and neurons are structurally unaltered in 3-month-old 5xFAD mice

Functional impairment in the NVC can result from structural alterations in the neurovascular unit. Previous reports have shown neurovascular structural changes in 5xFAD mice, including reduction in the vessel density, formation of the string vessels, neuronal loss and increased density of reactive astrocytes ^57,58^. We measured integrity of the neurovascular unit by analyzing density of capillaries and neurons in the cortex of control and 5xFAD mice. Capillary densities were measured using 3D-vessel *z-*stacks (350-400 μm depth from the pial surface) collected with plasma marker in the same FOVs, in which cECs Ca^2+^ activity was measured. Pial and penetrating vessels were removed during the analysis and only capillaries were used for the density calculations. Total capillary length was quantified for each z-stack and density was calculated as capillary length normalized to volume of the stack. Capillary densities were comparable between control and 5xFAD mice **(Supplement Figure 3G-H and Supplement movie S5-S6)** showing no alteration in the capillary density in 5xFAD mice. We next measured neuronal density in the coronal sections collected from the perfusion fixed mouse brain samples. We fluorescently labelled these coronal sections with neuronal marker, NeuN (**Figure 3O**) to quantify neuronal density. Cortical neuronal densities were comparable between control and 5xFAD mice (**Figure 3P),** indicating no detectable neuronal loss has occurred in 3-month-old 5xFAD mice. Collectively, these findings confirm that capillary and neuronal cells are structurally comparable between controls and 5xFAD mice.

### TRPV4 channel dysfunction leads to Electro-Calcium uncoupling and attenuates functional hyperemia in 5xFAD mice

We next evaluate the molecular mechanisms involved in the loss of cECs Ca^2+^ signals and in-turn crippled bimodal functional hyperemic response in 5xFAD mice. We recently showed cECs Ca^2+^ signals are elevated by membrane hyperpolarization, where this hyperpolarization increases driving force of Ca^2+^ entry into cECs through TRPV4 channels ^32^. This hyperpolarizing ‘electrical’ signal is generated via K_ir_2.1 channels and can propagate long-distance with a fast velocity (∼2 mm/s) ^26^. In addition to directly controlling CBF, these electrical signals increase driving force of Ca^2+^ entry for a slow sustained modulation of CBF ^32^. This intricate communication termed as E-Ca Coupling has kinetics that align well with the bimodal functional hyperemic response (**Figure 2**). We hypothesized that E-Ca coupling regulates the bimodal functional hyperemia, where the fast phase corresponds to the K_ir_2.1-induced electrical signals and the slow phase is regulated by Ca^2+^ signaling to maintain CBF during long-stimulation. Further disruption in E-Ca coupling alters fast-to-slow phase transition during functional hyperemia in 5xFAD mice. To test this hypothesis, we evaluated contributions of E-Ca coupling in control and 5xFAD mice by performing *in-vivo* cECs Ca^2+^ activity measurements in the presence of K_ir_2.1 channel blocker, Barium (Ba^2+^; 100 μM) (**Figure 4A-B**). Inhibition of K_ir_2.1 channels caused a significant reduction in cECs Ca^2+^ activity in the control mice confirming involvement of E-Ca coupling. However, Ba^2+^ had no effect on the cECs Ca^2+^ signals in 5xFAD mice showing a disruption in the E-Ca coupling (**Figure 4C**). Similarly, the number of Ca^2+^ events were significantly inhibited in the control mice in presence of Ba^2+^, but not in the 5xFAD mice (**Figure 4D**). This evidence supports an impairment in the E-Ca coupling in 5xFAD mice.

**Figure 4.**
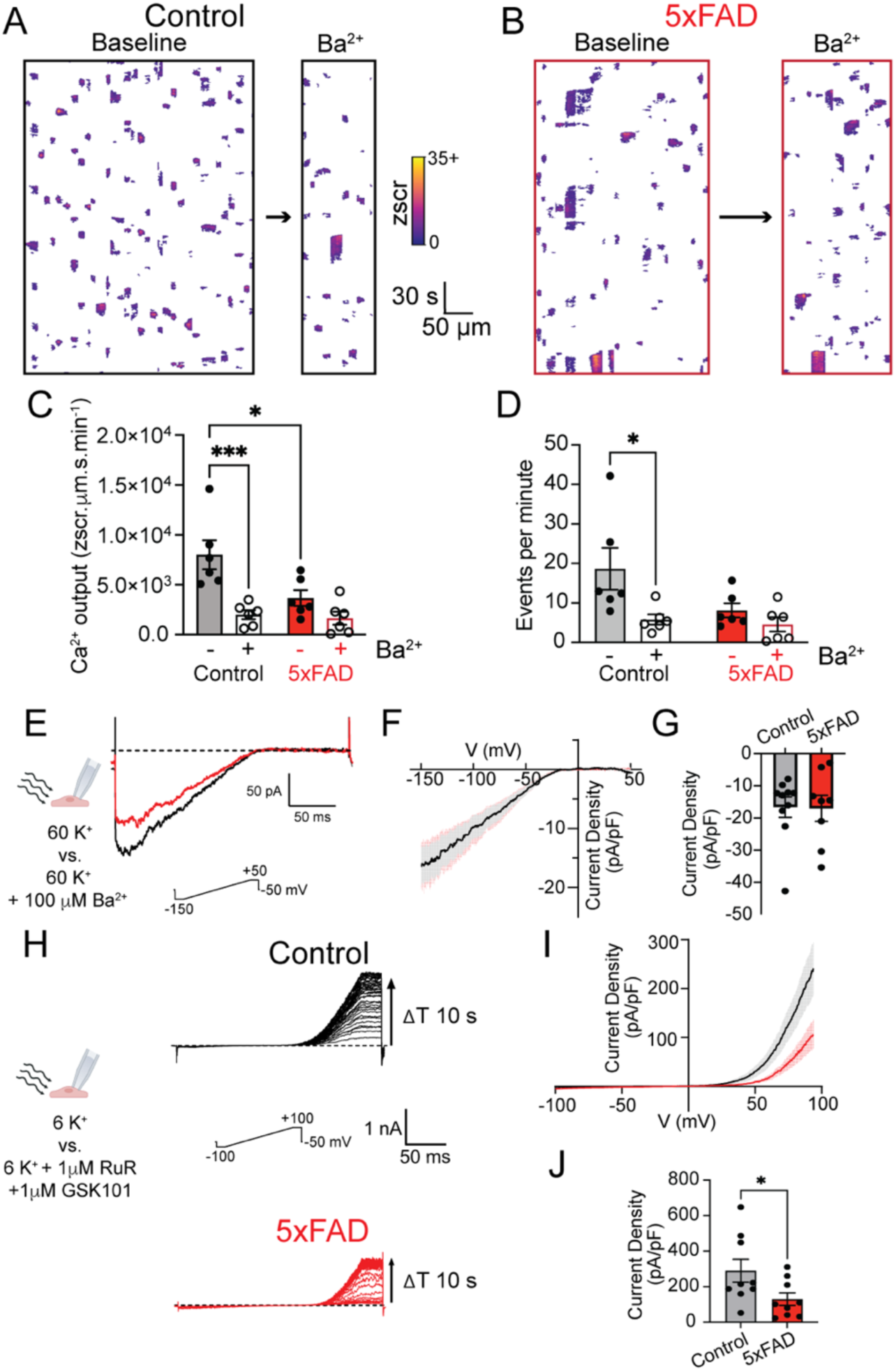
TRPV4 dysfunction leads to E-Ca uncoupling. **(A-B)** Spatiotemporal map of control **(A)** and 5xFAD **(B)** mice presenting cECs Ca^2+^ events without (left) or with (right) 100 µM Ba^2+^. **(C)** Comparison of cECs Ca^2+^ output between control and 5xFAD with (+) or without (−) 100 µM Ba^2+^ (Control-barium: 8014 ± 1450 Control+barium: 2014 ± 449, 5xFAD-barium: 3680 ± 780, 5xFAD+barium: 1657 ± 658 zscr·μm·s·min^-1^, Control-barium vs. Control+barium *** p<0.001, Control-barium vs. 5xFAD -barium * p<0.05, other comparisons ns, n=6, Two-way ANOVA with Tukey’s multiple comparison). **(D)** Comparison of cECs Ca^2+^ events between control and 5xFAD with (+) or without (−) 100 µM Ba^2+^ (Control-barium: 18.6 ± 5.3, Control + barium: 5.8 ± 1.3, 5xFAD-barium: 8.1 ± 1.8, 5xFAD-barium: 4.5 ± 1.8 events·min^-1^, control-barium vs control + barium, * p<0.05, other comparisons ns, n=6, Two-way ANOVA with Tukey’s multiple comparison). **(E)** *ex-vivo* patch clamp set-up for K_ir_2.1 channel whole cell current recordings, with 60 mM K^+^ bath with and without 100 µM Ba^2+^ (left). Representative Ba^2+^-sensitive current for control (black) and 5xFAD (red) is shown with voltage ramp protocol (−150 to +50 mV). **(F)** Averaged current density-voltage curve for controls and 5xFAD mice generated from ramp protocol with ramp velocity of 1 V/s. Note the identical conductance. **(G)** Comparison of steady-state current density of controls and 5xFAD mice at −150 mV (Control: −17 ± 3.2 vs. 5xFAD: −17 ± 4.1 pA/pF, ns, unpaired t-test, 8-10 cells from n=5 mice). **(H)** *ex-vivo* patch clamp set-up for TRPV4 channel currents recordings, with 6 mM K^+^ bath with and without 1 µM GSK101 and 1 µM RuR (left). Representative time course for GSK101-induced increase in TRPV4 currents in cECs isolated from control (black) and 5xFAD (red) mice shown with voltage ramp protocol (−100 to +100 mV). **(I)** Averaged current density-voltage curve for control and 5xFAD generated from ramp protocol with ramp velocity of 1 V/s. **(J)** Comparison of GSK101-saturated steady-state current density of control and 5xFAD at +100 mV (Control 290 ± 64 vs. 5xFAD: 130 ± 35 pA/pF, *p<0.05, unpaired t-test, 9 cells from n=4-5 mice).

To further investigate the deficiency in E-Ca coupling, we measured K_ir_2.1 and TRPV4 channel currents in freshly isolated cECs; two ion channels that are essential for E-Ca coupling ^32^. We measured K_ir_2.1 channel currents with conventional whole cell patch clamp recordings in 60 mM K^+^ bath solution with 200-ms voltage ramps from −150 to +50 mV and holding potential of −50 mV (**Figure 4E**), with high external K^+^ chosen to increase K_ir_2.1 channel conductance. Robust inward currents were detected at the voltage negative to the equilibrium potential for K^+^ (E_k_ ∼-23 mV) and these currents were eliminated upon application of 100 µM Ba^2+^, consistent with the previous reports ^26,59^. Ba^2+^-sensitive traces and current density were compared between 5xFAD mice and controls. Comparison of I-V curves between control and 5xFAD mice exhibited similar conductance with the steady-state current density of approximately −17 pA/pF at −150 mV (**Figure 4F and G**). These findings suggest the deficiency in E-Ca coupling does not arise from the reduction in K_ir_2.1 channel activity. Next, we evaluated whole-cell current density of TRPV4 channels by using 6 mM K^+^ bath solution with selective TRPV4 agonist GSK1016790A (hereafter GSK101; 1 µM) and a voltage-dependent pore blocker Ruthenium red (RuR, 1 µM), which blocks inward Ca^2+^ currents through TRPV4 channels. The currents were evoked using 200-ms voltage ramp from −100 to +100 mV applied every 10 second over the course of 10 min, allowing sufficient time for GSK101 to induce maximal TRPV4 currents (**Figure 4H**). Comparison of I-V curves showed that outward currents through TRPV4 channels were greatly reduced in 5xFAD compared to the controls despite the same recording conditions (**Figure 4I**), with the steady-state current density at +100 mV for 5xFAD (130 ± 35 pA/pF) significantly reduced compared to the controls (290 ± 64 pA/pF) (**Figure 4J**).

We next examined the mechanisms underlying the impairment of fast-to-slow phase transition of functional hyperemia using EC-specific K_ir_2.1 KO mice. We hypothesized that EC K_ir_2.1 KO mice would exhibit a prolonged fast phase compared to 5xFAD mice while showing a similar disrupted fast-to-slow phase transition due to impaired E-Ca coupling, as observed in 5xFAD mice. Consistent with this hypothesis, EC K_ir_2.1 KO mice had a significant reduction in whisker stimulation-induced hyperemia with marked impairment in fast and slow phases (**Figure 5A**). The steady state increase in CBV was significantly reduced in EC K_ir_2.1 KOs (CBV: 6.4 ± 1.5%) compared to controls (**Figure 5B**). Moreover, the fast phase was significantly prolonged in in EC K_ir_ 2.1 KOs (9.91 ± 4.5 s) compared to controls (2.40 ± 0.32 s) or 5xFAD mice (3.93 ± 0.46 s) (**Figure 5C-D and Figure 2F**). These findings align with our electrophysiology measurements showing comparable cECs K_ir_2.1 channel activity between controls and 5xFAD mice and confirm that loss of EC K_ir_2.1 channel activity is not solely responsible for crippled functional hyperemia in 5xFAD mice. Additionally, EC K_ir_2.1 KOs had a delayed fast-to-slow phase transition (ΔT_drop_) with a larger Amplitude drop (A_drop_) (**Figure 5E-F**) as observed response in 5xFAD mice. These findings reinforce the idea that the E-Ca coupling impairment can result from reduced activity of either Kir2.1 or TRPV4 channels, however in 5xFAD mice, these early impairments are due to functional loss of TRPV4 channels. Collectively, the reduction in TRPV4 channel activity with unaltered K_ir_2.1 channel activity in 5xFAD mice confirms that impaired Ca^2+^ influx through TRPV4 channels disrupts E-Ca coupling, corresponds to a crippled fast-to-slow phase transition in the functional hyperemia.

**Figure 5.**
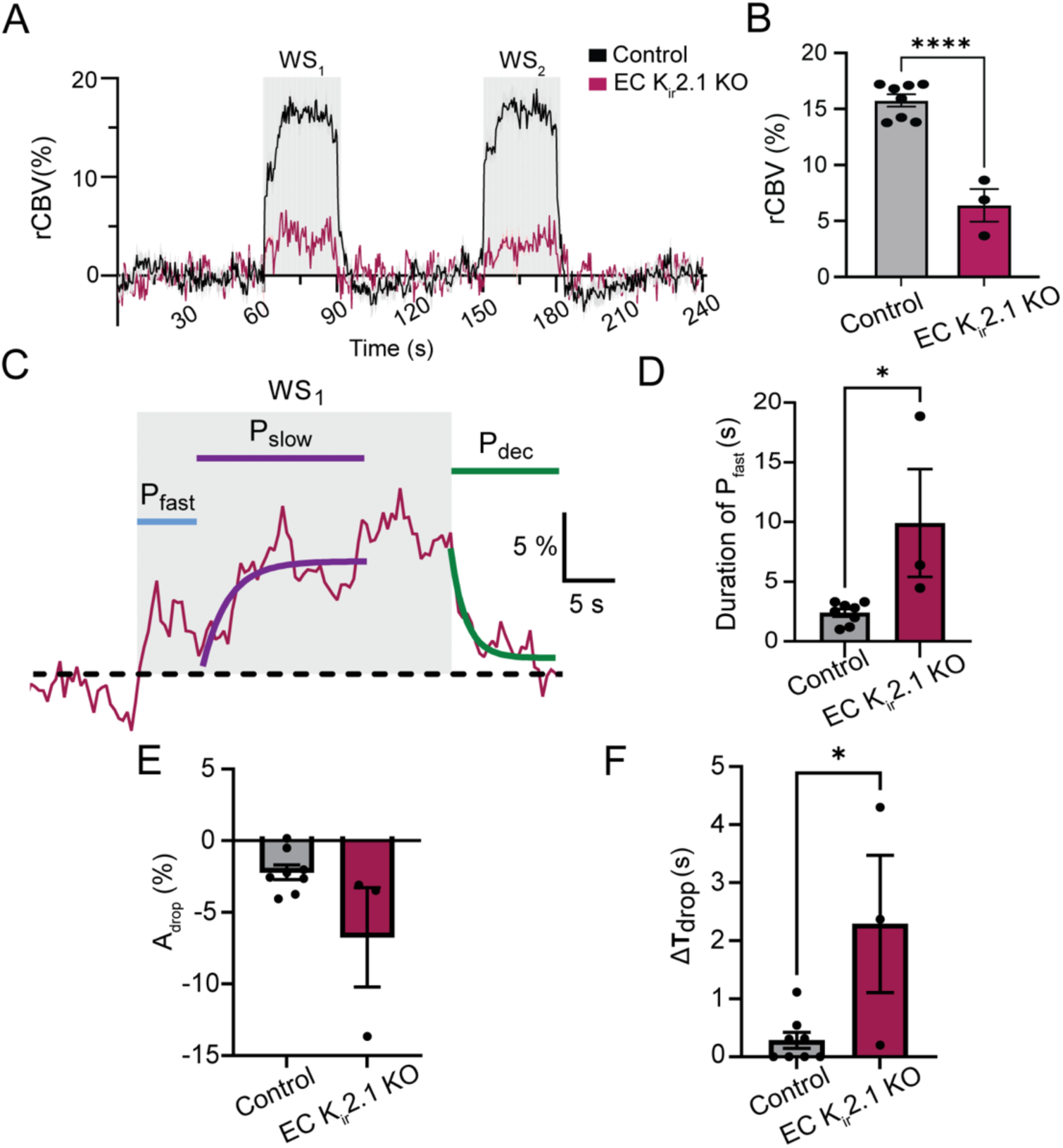
Fast phase and the fast-to-slow phase transition of functional hyperemia are impaired in EC K_ir_ 2.1 KO mice. **(A)** Averaged time course of normalized rCBV change in control (black, same as in Fig.2B) and EC K_ir_2.1 KO (maroon), with same recording parameters as in Fig.2B. **(B)** Comparison of steady-state change in rCBV at WS_1_ between controls and EC K_ir_2.1 KOs. The EC K_ir_2.1 KO mice show reduction in steady-state rCBV change (Control: 16 ± 0.55, n=8 vs. EC K_ir_2.1 KOs: 6.4 ± 1.5 %, **** p<0.0001, unpaired t-test, n=3). **(C)** Representative trace of EC K_ir_2.1 KO showing WS_1_ period. **(D)** Comparison of fast phase duration between control and EC K_ir_2.1 KO mice (Control: 2.4 ± 0.32 vs EC K_ir_2.1 KO: 9.9 ± 4.5 sec, * p<0.05, n=3-8).**(E-F)** Comparison of two transition impairment parameters: amplitude drop (A_drop,_ Control: −2.2 ± 0.51 vs. EC K_ir_2.1 KO: −6.7 ± 3.5 %, ns, unpaired t-test, n=3-8) and difference in transition time between fast and slow phase (ΔT_drop_, Control: 0.29 ± 0.14 vs. EC K_ir_2.1 KO:2.3 ± 1.2 sec, *p<0.05, unpaired t-test, n=3-8).

## 4. DISCUSSION

We demonstrate that 5xFAD mice have neurovascular uncoupling as an early neurovascular pathological event during the disease development. These neurovascular deficits impair functional hyperemia and precede neuronal loss or dysfunction, common hallmarks of AD. Our functional studies confirmed that neurovascular deficits can be attributed to the impairment in E-Ca coupling with a robust reduction in cECs TRPV4 channel activity (**Figure 6**). Despite these functional impairments, capillary structure and resting cerebral perfusion remain unaltered. To our knowledge, cECs E-Ca uncoupling described here could be the earliest vascular changes in AD pathology preceding the neurodegeneration and long-term cognitive decline.

**Figure 6.**
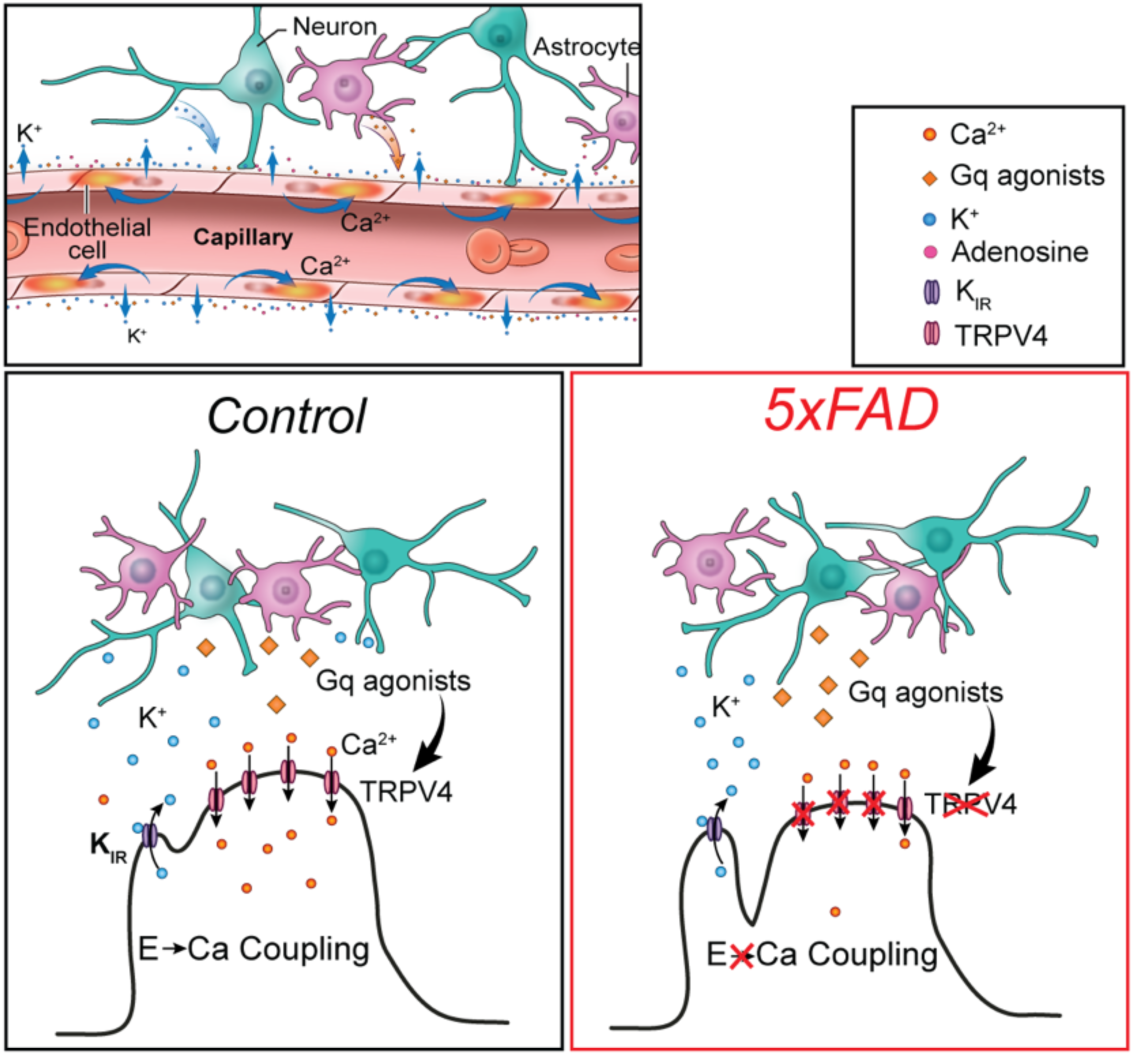
Graphical abstract showing E-Ca coupling (top) in physiology. Contribution of E-Ca coupling in functional hyperemia in Control mice (bottom left) and crippled E-Ca coupling in 5xFAD mice (bottom right).

The brain vascular system supplies essential nutrients and clears metabolic byproducts generated with neuroglial activity. Although the human brain accounts for 2% of body weight, it consumes ∼one fifth of the body’s energy, where maximum energy is consumed in the action potentials and postsynaptic glutamate activity ^5,6^. Imbalance in the neurovascular unit integrity—whether structural or functional—contribute to the impaired cerebral hemodynamics reported in AD patients as early as the presymptomatic or MCI phase. Early cerebral hemodynamic changes are observed both in the familial and sporadic late-onset AD patients, as well as in the corresponding animal models^60,61^. These findings prompted us to investigate the neurovascular deficits that emerge during the presymptomatic phase of AD using the 5xFAD mouse model. Identification of early neurovascular deficits informed us about the contributions of vascular pathophysiological changes in the initial progression of the disease.

Mild AD patients reportedly have deficiency in functional hyperemia and BOLD signals when given visual stimulations, and these deficits become more severe with the age ^62,63^. Furthermore, mild AD patients have reduced cortical glucose consumption and metabolic rate^64^. Despite these impairments in the functional hyperemia and brain energy consumption, mild AD patients have preserved resting cerebral perfusion ^62,63^. Likewise, 4-month-old 5xFAD mice are reported to have preserved baseline perfusion with a trend towards reduction around 7-months^65^. In contrast to this, another study reported no change in the resting perfusion up to 12-month of age in 5xFAD mice ^66^. Our previous work showed a reduction in the whisker stimulation induced functional hyperemia in 12-month-old 5xFAD mice ^11^. Follow up work by Silva et al. reported impaired functional hyperemia in 5-6 months-old mice ^12^. These studies are some examples showing quantifiable deficits in the functional hyperemia in the symptomatic phase ^11,12,65,67^. However, little is known about the neurovascular deficits involved in cerebral hemodynamics changes during the presymptomatic stage. In this study, we chose 3-month-old 5xFAD mice that did not have measurable behavioral deficits in the spatial learning and memory. These 5xFAD mice showed a robust reduction in whisker stimulation-induced functional hyperemia, while their resting-state cerebral perfusion was preserved. The functional hyperemia showed a bimodal response consisting of two distinct phases, namely fast and slow with sustained neural activity. Both controls and 5xFAD mice presented the bimodal functional hyperemic response, however the temporal kinetics were different. The initial fast phase was largely preserved in the 5xFAD mice, while fast-to-slow phase transition was significantly impaired. The bimodal functional hyperemic response relies on multiple parallel NVC mechanisms in concert ^52,53,68^. Previous work has described that the fast phase is governed by neuronal mediators, while the slow phase relies on the astrocytic mediators ^52^, however concomitant vascular signaling mechanisms involved in each phase remains unknown.

We recently described E-Ca coupling that involves close communication between two distinct NVC mechanisms; electrical and Ca^2+^ signaling^32^. The spatiotemporal characteristics of E-Ca coupling align well with the bimodal functional hyperemic response, where electrical signals could contribute to initial ‘fast’ phase and Ca^2+^ signaling maintains CBF in the ‘slow’ phase with sustained neural stimulation. The electrical component in the E-Ca coupling involves rapid membrane hyperpolarization of the cECs mediated by K_ir_2.1 channels ^26,32^. K_ir_2.1 channel activity in cECs was preserved in 5xFAD mice. Moreover, the kinetics of fast phase of functional hyperemia were significantly different between EC-Kir2.1 KOs and 5xFAD mice. These findings confirmed that electrical component of the E-Ca coupling corresponds to the fast phase of functional hyperemia and are possibly unaltered in 3-month-old 5xFAD mice. In contrast, Ca^2+^ signaling of the E-Ca coupling is impaired with reduced TRPV4 channel activity. Dysfunction of TRPV4 channels may exert a ‘double hit’ effect on functional hyperemia. First, impaired TRPV4 channels may manifest as a delay in the transition from the fast-to-slow phase. Second, the reduced magnitude Ca^2+^ influx through TRPV4 channels is likely to compromise subsequent Ca^2+^ induced Ca^2+^ release (CICR) from intracellular ER stores, resulting in an overall attenuated response in slow phase. These mechanistic insights about impaired functional hyperemia due to TRPV4 channel dysfunction and crippled E-Ca coupling could be one of the earliest changes in 5xFAD mice and can precede impairment in EC K_ir_2.1 channel functions^11^. These findings may direct future studies for therapeutic interventions to improve CBF response in AD.

Clinical studies report mixed findings regarding cortical capillary length in AD patients, with the variability likely reflecting differences in the disease stage, patient demographics and experimental methodologies ^69–72^. The influence of disease stage is illustrated in the retina microvasculature where microvascular density remains unchanged in the patients with mild cognitive impairment (MCI) but is significantly reduced in the AD patients ^73^. Similar reduction in the cortical microvascular density is reported in the animal models of AD including 5xFAD mice ^69–72^. Longitudinal studies evaluating cortical cerebrovascular changes in 5xFAD mice have reported decreased vessel length and microvascular damage beginning around 4-months of age ^58,74^. These changes in the vascular density can contribute to impaired functional responses including functional hyperemia and E-Ca coupling. However, we did not observe any changes in cortical capillary density in 3-month-old 5xFAD mice, suggesting capillary structural changes are not contributing to the observed neurovascular deficits.

Neuronal activity is the essential driver for NVC including E-Ca coupling ^32^. A decrease in the neural mediators, *e.g.,* K^+^, G_q_ agonists, and ROS can also cripple NVC. We evaluated neuronal density and functions, where both were preserved in 5xFAD mice. Clinical studies suggest a biphasic trajectory of neuronal activity in AD, with early cortical and hippocampal neuronal hyperexcitability in MCI phase followed by hypoactivity in the later disease stages ^75–77^. We also observed subtle hyperactivity in the somatosensory neurons of 5xFAD mice, characterized by high frequency of large-amplitude events. However, these changes were modest compared to pronounced reduction in capillary vascular signaling associated with E-Ca uncoupling and loss of TRPV4 channel activity. Together, these findings support the notion that neurovascular deficits precede the neurodegeneration in 5xFAD mice. These early deficits may contribute to the cognitive decline in 5xFAD mice, as mild cortex-dependent memory impairments are reported at ∼4 months progressing to more pronounced deficits by 4–6 months ^78–83^. Follow up studies aiming to restore TRPV4 functional deficits and in-turn E-Ca coupling—alongside assessment of cognitive functions— may provide insight into potential therapeutic interventions for AD.

Altogether, this study includes multiple complementary approaches that collectively reinforce the conclusion that early neurovascular deficits precede neurodegeneration and cognitive impairment in 5xFAD mice. Nonetheless, a few limitations should be considered. 5xFAD model develops amyloid-β plaque deposition as early as 1.5–2 months of age ^84^, in conjunction with increased levels of soluble amyloid-β. Although this study identifies early molecular mechanisms of neurovascular uncoupling in 5xFAD mice, the influence of soluble amyloid-β on the observed NVC deficits cannot be excluded as the pre-plaque mechanisms. Moreover, our conclusions are derived from male mice and cannot exclude the sex-based differences. In addition, the attenuation of the slow-phase vascular response observed here was not accompanied by direct assessment of astrocytic function. Given the role of astrocytes in NVC, altered astrocytic signaling may contribute to this deficit and should be addressed in future studies.

In conclusion, we provide molecular insights for impairment in functional hyperemia during early phase of AD. These early neurovascular deficits coincide with subtle neuronal hyperexcitability. Given the shared pathophysiology of familial AD with sporadic AD and cerebral amyloid angiopathy (CAA), our findings may also inform studies of early neurovascular deficits in sporadic AD and CAA. Importantly, our results suggest that sensory-evoked CBF changes are early hallmarks of familial AD pathophysiology, preceding alterations in the resting cerebral perfusion and neuronal loss. Together, these findings reveal early neurovascular uncoupling in AD that may be leveraged for early detection and therapeutic interventions.

## Supplement movie legends

Movie S1. Capillary endothelial Ca^2+^ activity in the cortical layer II-III of a control mouse.

Movie S2. Capillary endothelial Ca^2+^ activity in the cortical layer II-III of a 5xFAD mouse.

Movie S3. Widefield cortical neuronal Ca^2+^ signals from a control mouse.

Movie S4. Widefield cortical neuronal Ca^2+^ signals from a 5xFAD mouse.

Movie S5. 3D-rendering showing the cortical capillary distribution from a control mouse.

Movie S6. 3D-rendering showing the cortical capillary distribution from a 5xFAD mouse.

## Acknowledgements

We acknowledge and thank T. Byrd for her technical assistance; Dr. G. W. Hennig (University of Vermont), Dr. M. Weston (Virginia Tech) and E. Jin Choi for their advice in the development of imaging analysis tools; Dr. J. Munasinghe, Dr. S. Dodd and Dr. S. Yadav (John Hopkins University) for technical advice with fMRI experiments; Dr. S. Bradley and Dr. Y. Chudasma for their guidance in the behavioral experiments as this work was supported, in part, by the NIMH IRP Rodent Behavioral Core (MH002952); NIMH Section of Instrumentation staff for their help in custom fabrication and controllers for image acquisition; NICHD Histology core for technical assistance with immunofluorescence imaging; Dr. M. Kotlikoff (Cornell University) for providing *Cdh5*-GCaMP8 mice. Some of the schematic figures were created with BioRender. *Smith, J. (2025). BioRender.com/c248457.* We utilized ChatGPT (https://www.chatgpt.com) for language editing assistance. After using, the author(s) reviewed and edited the content as needed and take(s) full responsibility for the content of the published article.

## Author contribution

A.M. conceptualized and supervised the project; A.M. and H.C.K. wrote the initial draft of the manuscript; H.C.K., A.E., M.M. performed experiments, analyzed results and prepared the figures; J.L., J.G., S.H., Y.L. developed image analysis tools and analyzed data; H.C.K., A.E., M.M., J.L., J.G, S.H, Y.L., M.T.N., A.K. and A.M. reviewed and edited the manuscript.

